# POLARIS is a copper-binding peptide required for ethylene signalling control in *Arabidopsis*

**DOI:** 10.1101/2023.06.15.545071

**Authors:** Anna J. Mudge, Saher Mehdi, Will Michaels, Beatriz Orosa-Puente, Weiran Shen, Charlie Tomlinson, Wenbin Wei, Claudia Hoppen, Buket Uzun, Dipan Roy, Flora M. Hetherington, Jennifer F. Topping, Ari Sadanandom, Georg Groth, Nigel J. Robinson, Keith Lindsey

## Abstract

Ethylene signalling represents one of the classic hormonal pathways in plants, with diverse roles in development and stress responses. The dimeric ethylene receptor localizes to the endoplasmic reticulum (ER) and contains Cu(I) ions essential for ethylene binding and signal transduction. We previously discovered that mutants in the *Arabidopsis* gene *POLARIS* (*PLS*), encoding a 36 amino acid peptide, exhibit enhanced ethylene signalling responses, suggestive of reduced receptor activity, but the role and activity of the peptide in this signalling cascade has not been defined. Here we report PLS binds copper as a 1:2 thiol-dependent Cu(I):PLS_2_ complex, with an affinity of 3.79 (±1.5) x10^19^ M^-2^, via two cysteine residues also found in the related species *Camelina sativa*. These residues are also essential for biological function. This affinity precludes PLS as a cytosolic Cu chaperone. We demonstrate that PLS localizes to endomembranes and interacts with the transmembrane domain of receptor protein ETR1. PLS-ETR1 binding is increased in the presence of copper, and this interaction provides a Cu-dependent mechanism for mediating a repression of ethylene responses. *PLS* transcription is up-regulated by auxin and down-regulated by ethylene, and so PLS-ETR1 interactions also provide a mechanism to modulate ethylene responses in high auxin tissues.

## INTRODUCTION

Ethylene is used by plants as a gaseous hormone to regulate many aspects of development and responses to biotic and abiotic stresses (Johnson and Ecker, 1998). It is perceived by a family of receptors that, in *Arabidopsis*, comprises 5 members, ETR1 (ETHYLENE RESPONSE 1), ERS1 (ETHYLENE RESPONSE SENSOR 1), ERS2, ETR2, and EIN4 (ETHYLENE-INSENSITIVE 4) (Chang et al, 1993; Hua et al., 1995, 1998; Sakai et al., 1998) located on the endoplasmic reticulum (ER) (Chen et al., 2002; Grefen et al., 2008). The receptors are related to bacterial two-component systems (Chen et al., 2002), form dimers through disulphide bonding at the N-terminal hydrophobic domains (Schaller et al., 1995; Hall et al., 2000) and contain Cu(I) ions bound to residues Cys65 and His69, essential for ethylene binding and signal transduction (Rodriguez et al., 1999; McDaniel and Binder, 2012). In the absence of ethylene these receptors activate a negative regulator CTR1 (CONSTITUTIVE TRIPLE RESPONSE 1), which is a mitogen-activated protein kinase kinase kinase (MAPKKK), so preventing ethylene responses (Chang, 2003; Gao et al., 2003). Mechanisms by which receptor activity is regulated are not fully understood.

Introduction of copper to the ER and ethylene receptor requires the RAN1 (RESPONSIVE TO ANTAGONIST1) protein. This is a predicted copper-transporting P-type ATPase homologous to the yeast Ccc2p and to human Menkes and Wilson disease proteins (Hirayama et al., 1999). Strong loss-of-function mutants of *RAN1* in *Arabidopsis* (e.g. *ran1-3*, *ran1-4*) exhibit an enhanced ethylene signalling response (Binder et al., 2010), consistent with a loss of receptor function, and similar to higher order loss-of-function receptor mutants, which also show an ethylene hypersignalling phenotype (Qu et al., 2007). Mechanisms of copper homeostasis at ETR1 are unknown, as is true for other compartmentalized cuproproteins supplied with copper, for example via Ccc2p, Menkes or Wilson ATPases. RAN1 in Arabidopsis localizes to endomembrane systems, including *trans*-Golgi and ER compartments, and is necessary for both the biogenesis of ethylene receptors and copper homeostasis - loss of function *ran1* mutants suggest copper is required for both ethylene binding and receptor function (Binder et al., 2010). RAN1 can interact directly with ETR1 and the copper chaperones ANTIOXIDANT1 (ATX1) and COPPER CHAPERONE (CCH), suggestive of a transfer of copper between proteins to deliver it to the ethylene receptor as part of the receptor biogenesis pathway at the ER (Hoppen et al., 2019).

Our understanding of receptor function is still incomplete, however. For example, how is Cu(I) delivery from RAN1 to ETR1 mediated? How does Cu(I) influence receptor conformation and function? Are there other Cu(I)-binding components involved? Is the receptor regulated in a tissue-specific way or in response to the hormonal environment in a tissue? Is this process part of the crosstalk mechanism with other hormone signaling pathways? It is well established that ethylene signaling interacts with and is impacted by other hormonal pathways but does this influence the receptor metalation state and with developmental consequences?

We have previously shown that the loss-of-function *polaris* (*pls*) mutant has some phenotypic similarities to *ran1* loss-of-function alleles and to *ctr1*, exhibiting a triple response phenotype (short hypocotyl and root, exaggerated apical hook, radial expansion) in the dark in the absence of ethylene (Chilley et al., 2006), and a short root in light-grown seedlings, consistent with its known expression in the root meristem in light-grown seedlings (Figure 1A). Transgenic complementation of the mutant by the *PLS* gene (AT4G39403), which encodes a 36 amino acids peptide, suppresses the mutant phenotype (Casson et al., 2002). The *pls* mutant phenotype is rescued by the gain-of-function ethylene resistant mutation *etr1-1* and pharmacological inhibition of ethylene signalling by silver ions (Chilley et al., 2006). Ethylene gas production in the *pls* mutant is at wild type levels, indicating the peptide plays a role in ethylene signalling rather than biosynthesis (Chilley et al., 2006). *PLS* transgenic overexpressers (PLSOx seedlings) in contrast exhibit a suppression of the triple response phenotype when grown in the presence of the ethylene precursor ACC, similar to the gain of function *etr1-1* mutant, but this suppression is incomplete (PLSOx seedlings show some response to ACC; Casson et al., 2002; Chilley et al., 2006). PLS overexpression also partially suppresses the *ctr1* mutant phenotype, indicating that the PLS peptide acts upstream of CTR1 (Chilley et al., 2006).

**Figure 1.**
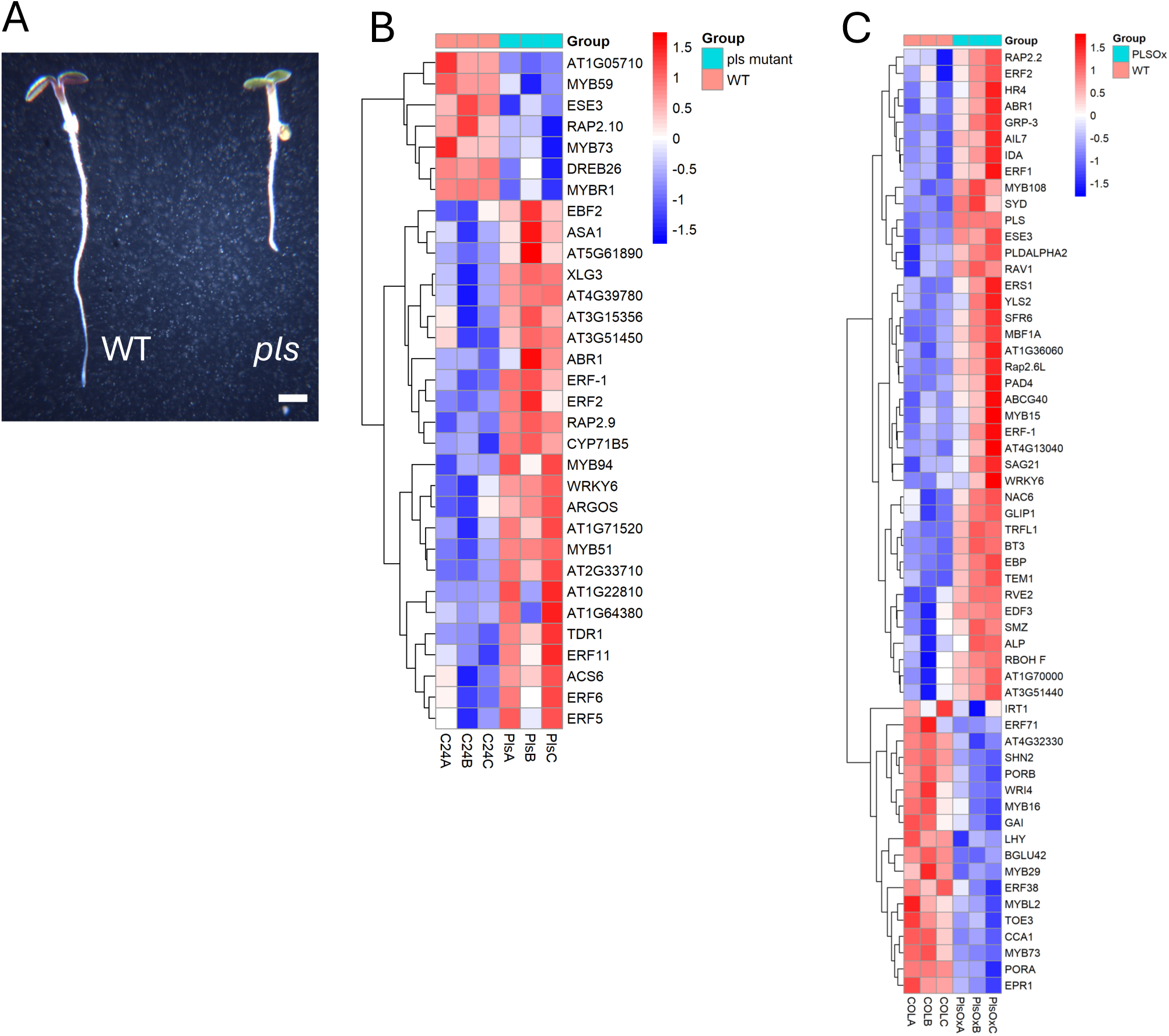
The PLS peptide is required for ethylene control of seedling growth. (**A**). Wild type (left) and *pls* mutant (right); bar = 5 mm. (**B-C**) Heat maps showing expression levels of 32 ethylene-responsive genes in *pls* (**B**) and of 58 genes in *PLS* overexpressing (PLSox) seedlings (**C**), compared to wild type levels. Data for 3 biological replicates are shown and are expressed as log_2_-fold changes in *pls* mutant and PLSox seedlings compared wildtype with respective significance levels (*P-adj* value) provided in Tables S1 and S2. A *P* < 0.05 and log_2_-fold of change of ± 0.5 was chosen to identify differentially expressed genes.

In this paper we show that the POLARIS peptide is required for correct ethylene responses, binds copper via two cysteine residues essential for biological function, but is unlikely to act as a metallochaperone intermediary; however it co-localises, and forms a copper-adduct with, ETR1. We suggest this provides cell type control and a crosstalk regulatory mechanism for ethylene responses through a role in its physical interaction with the ethylene receptor.

## RESULTS

### The POLARIS peptide is a negative regulator of ethylene responses

The PLS peptide is translated from a low-abundance transcript that, in light-grown seedlings, is most strongly expressed in the embryo and silique, seedling root tip and leaf vascular tissues (Figure S1), a pattern reflected in promoter-reporter expression patterns (Casson et al., 2002). To further investigate the *pls* molecular phenotype, we carried out an RNA-seq analysis on loss-of-function *pls* mutant and PLSOx transgenic overexpressor seedlings to identify differentially regulated genes, with mutant and overexpressor each compared with wild type seedlings of the same ecotype as controls (Fig. 1B,C). The *pls* mutant expresses no full length PLS coding sequence due to a T-DNA insertion in the coding region of *PLS* and disrupting function (Casson et al., 2002), while the PLSOx seedlings express significantly higher levels of *PLS* transcript compared to wild type (log_2_ fold 10.8, *padj* value = 2.02 x 10^-33^; Tables S1, S2). 836 genes were significantly up-regulated and 292 down-regulated in the *pls* mutant compared to wild type control seedlings (padj value <0.05) (Table S1). 1487 genes were significantly up-regulated and 1281 down-regulated in PLSOx seedlings compared to wild type (padj value <0.05; Table S2). Gene Ontology (GO) analysis of genes upregulated in *pls* mutant seedlings compared to wild type showed significant enrichment of genes associated with responses to hormone signalling, biotic and abiotic defence responses, and cell death (Table S3; Figure S2A).

Out of 307 GO:0009723 (response to ethylene) genes, 25 were significantly up-regulated and 7 down-regulated in the *pls* mutant compared with the wild type, with 40 up-regulated and 18 down-regulated in the PLSOx seedlings (Figure 1C; Table S3, S4, S5 and S6; Mudge et al., 2024), indicating a requirement for control over *PLS* expression levels for correct ethylene responses. While GO:0009723 (response to ethylene) is significantly enriched in genes up-regulated in the *pls* mutant compared to wild type (*FDR = 0.00062*; Table S3), a large number of up-regulated genes in *pls* are significantly associated with immunity, response to pathogens and hypersensitive response (Table S3 and Figure S2A). Down-regulated genes in *pls* were significantly enriched in GO categories of hormone biosynthetic processes (GO:0042446, *FDR = 0.038*), hormone metabolic process (GO:0042445, *FDR = 0.0096*) and regulation of hormone levels (GO:0010817, *FDR = 0.0013*) (Table S4, Figure S2B; Mudge et al., 2024), consistent with our previous studies that described PLS-dependent crosstalk between ethylene, auxin and cytokinin signalling (Liu et al., 2010, 2013; Moore et al., 2015). In PLSOx overexpressors, both up-and down-regulated genes similarly included those associated with GO terms hormone response, stress response and immune response, and genes associated with photosynthesis are downregulated in PLSOx seedlings, consistent with repressed photosynthetic activity in root tips (Table S5 and S6, Figure S2C, D; Casson et al., 2002). These data are consistent with a role for PLS in a number of ethylene responses and other related signalling, stress and developmental processes.

To understand better the relationship between PLS peptide structure and function between species, we carried out hydroponic feeding experiments using synthetic versions of PLS peptide from both *Arabidopsis* and its close relative *Camelina sativa*. *C. sativa* contains a gene with partial sequence identity to the *Arabidopsis PLS* gene, encoding a predicted peptide sequence that is 22 amino acids long and identical to the N-terminal 22 amino acids of the *Arabidopsis* PLS except for a phenylalanine to serine substitution at position nine (Figure 2A). We synthesized full-length PLS peptide, PLS(FL) and truncated versions from both *Arabidopsis* and *C. sativa* (Figure 1A) and supplied the peptides to *Arabidopsis pls* mutant seedlings hydroponically. The full-length peptides from both *Arabidopsis* and *C. sativa* and the N-terminal 22 amino acids sequence of the *Arabidopsis* peptide (N1) were each able to rescue the short primary root length of the *Arabidopsis pls* mutant (Figure 2B), similar to transgenic overexpression and genetic complementation using the wild type *PLS* coding sequence (Casson et al., 2002; Chilley et al., 2006). The mean primary root length of wild type (C24) seedlings did not change across all peptide treatments, while supply of *pls* seedlings with 50 nM PLS(FL) and N1 peptides showed a rescue of root growth (Figure 2C), and *t*-tests showed there was no significant difference between the rescue effects of PLS(FL) and PLS(N1); *P* > 0.1, n = 25). However, neither a 9 amino acids sequence (N2, Figure 2C) from the N-terminus, nor C-terminal sequences of 14 (C1) or 24 (C2) amino acids from *Arabidopsis* PLS were able to rescue the mutant (Figure 2C); each of these shorter peptides lacks one of the two Cys residues found in the functional longer length peptides PLS(FL) and PLS(N1). A fluorescent tagged (5-carboxyfluorescein, 5-FAM) version of the *Arabidopsis* N-terminal 22 amino acids sequence of the peptide (N1) is taken up by the roots, and also rescues the mutant root phenotype (Figures S3A, B).

**Figure 2.**
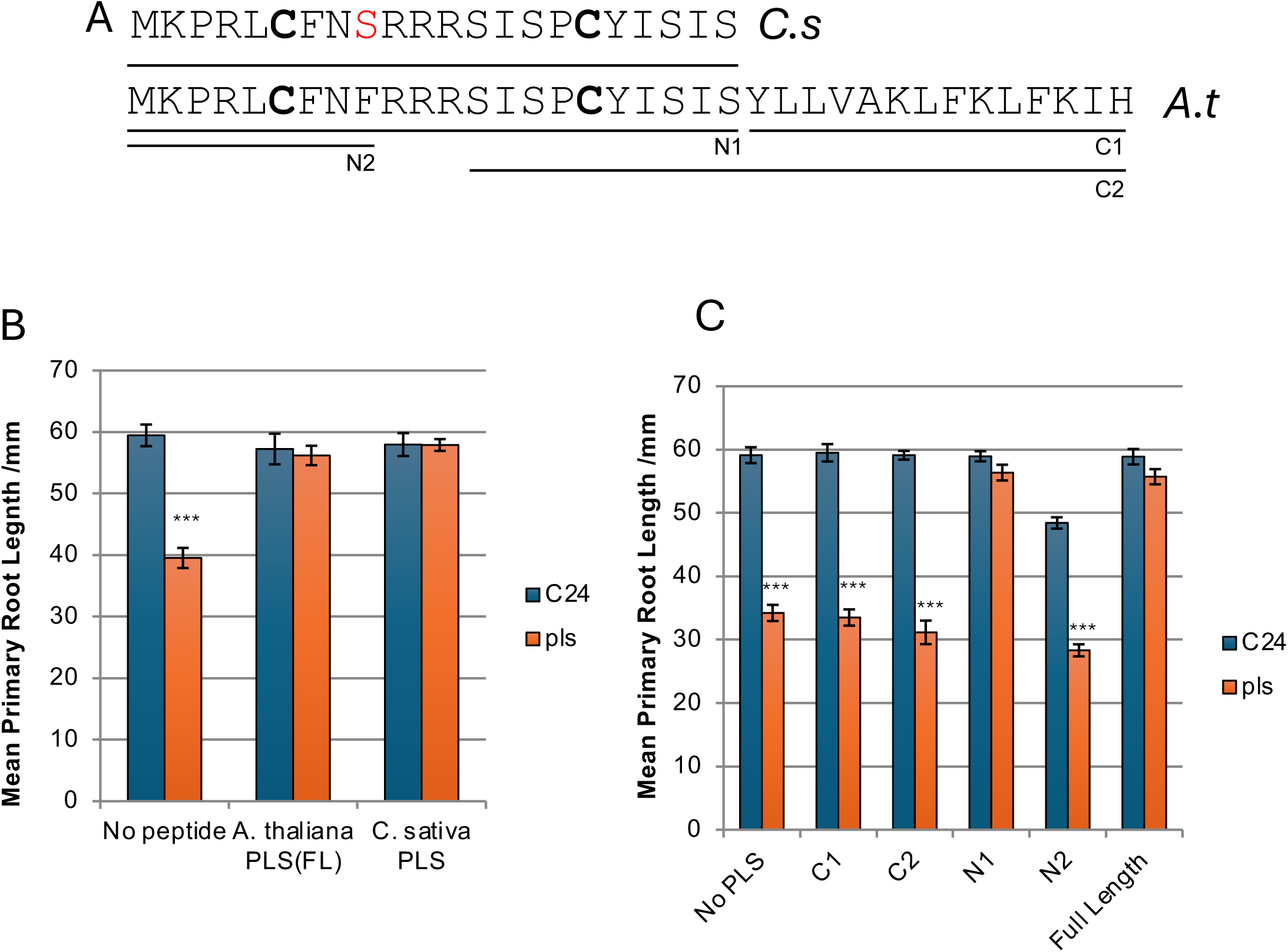
The PLS peptide is structurally and functionally conserved and complements the *Arabidopsis pls* mutant. (**A**). Amino acid sequence of the PLS peptide from *Arabidopsis thaliana* (*A.t*) with *Camelina sativa* PLS sequence (*C.s*), and synthetic truncations N1, N2, C1 and C2, indicated by horizontal lines. Two cysteine residues are highlighted in bold. (**B**) Effect of *Arabidopsis* PLS full length peptide, *A.t.* PLS(FL), and *Camelina* PLS peptide (*C.s.* PLS) on *Arabidopsis* primary root length. Wildtype (C24; blue bars) and *pls* mutant seedlings (red bars) were grown hydroponically in the presence (100 nM) or absence of peptide for 10 d. ***: *P* <0.05, *t*-test between C24 and *pls*, n = 20-25. (**C**) Effect of PLS full length and truncated peptides on wild type (blue bars) and *pls* mutant (red bars) *Arabidopsis* primary root length. Seedlings were grown hydroponically in the presence of 50 nM peptide for 10 d. C1 = C-terminal 14 amino acids, C2 = C-terminal 24 amino acids, N1 = N-terminal 22 amino acids, N2 = N-terminal 9 amino acids, full length PLS = 36 amino acids. Error bars show ± 1 standard error, n = 25. ***: *P* <0.05, *t*-test between C24 and *pls*, n = 25.

### PLS localizes to endomembranes including the endoplasmic reticulum

Since genetic studies suggest that PLS acts close to the ethylene receptor (Chilley et al., 2006), we hypothesized that it should localize to the same subcellular compartment as ETR1. The ethylene receptor in *Arabidopsis* is localized predominantly to the ER (Chang, 2003), and a *proPLS::PLS:GFP*-generated fusion protein (PLS:GFP) was used to investigate sub-cellular localization. Of five independent transformants with a single-copy insertion of the *proPLS::PLS:GFP* gene in a *pls* mutant background, four were found to complement fully the *pls* mutant (Figure S4), similar to the wild type cDNA (Casson et al., 2002), demonstrating functionality of the gene fusion. The PLS:GFP was found to co-localize in transgenic plants with the ER marker dye ER-Tracker red^TM^, which binds sulphonylurea receptors of ATP-sensitive channels on the ER (Thermofisher; Figures 3A-C). It also co-localizes with an ER lumen-targeted red fluorescent protein RFP-HDEL (Lee et al., 2013) (Figures 3G-I). These observations suggest PLS:GFP locates both on the ER membrane and in the lumen. PLS:GFP also appears to localize to the nucleus and cytoplasm (Figure 3). As controls, free GFP protein expressed under the control of the *PLS* promoter is not co-localized to the ER (Figures 3D-F) and, as expected, the Golgi marker SH:GFP does not co-localize with ER Tracker (Figures 3M-O). *Trans*-Golgi-localized SULFOTRANSFERASE1 (ST1) mCherry (Bauer and Papenbrock, 2002) visualization shows PLS:GFP does not localize to the Golgi (Figures 2J-L). To further clarify the side of the ER membrane on which PLS localizes, transient expression of redox-sensitive GFP (roGFP2) fusions of PLS were carried out. The different excitation properties of roGFP2 in an oxidizing (ER lumen) or reducing environment (cytosol) allows discrimination of the precise location of PLS fused to RoGFP2. Ratiometric analysis and comparison with proteins of known localization (i.e. cytosolic RoGFP2, ER luminal RoGFP2, cytosolic v-SNARE SEC22-GFP, ER luminal RoGFP2-SEC22; Brach et al., 2009) revealed that PLS as either an N-or C-terminal roGFP2 fusion resides at the cytosolic side of the ER (as well as to other vesicular compartments; Figure 3P). However, there may be some translocation of PLS:GFP into the ER lumen as indicated by co-localization with RFP:HDEL.

**Figure 3.**
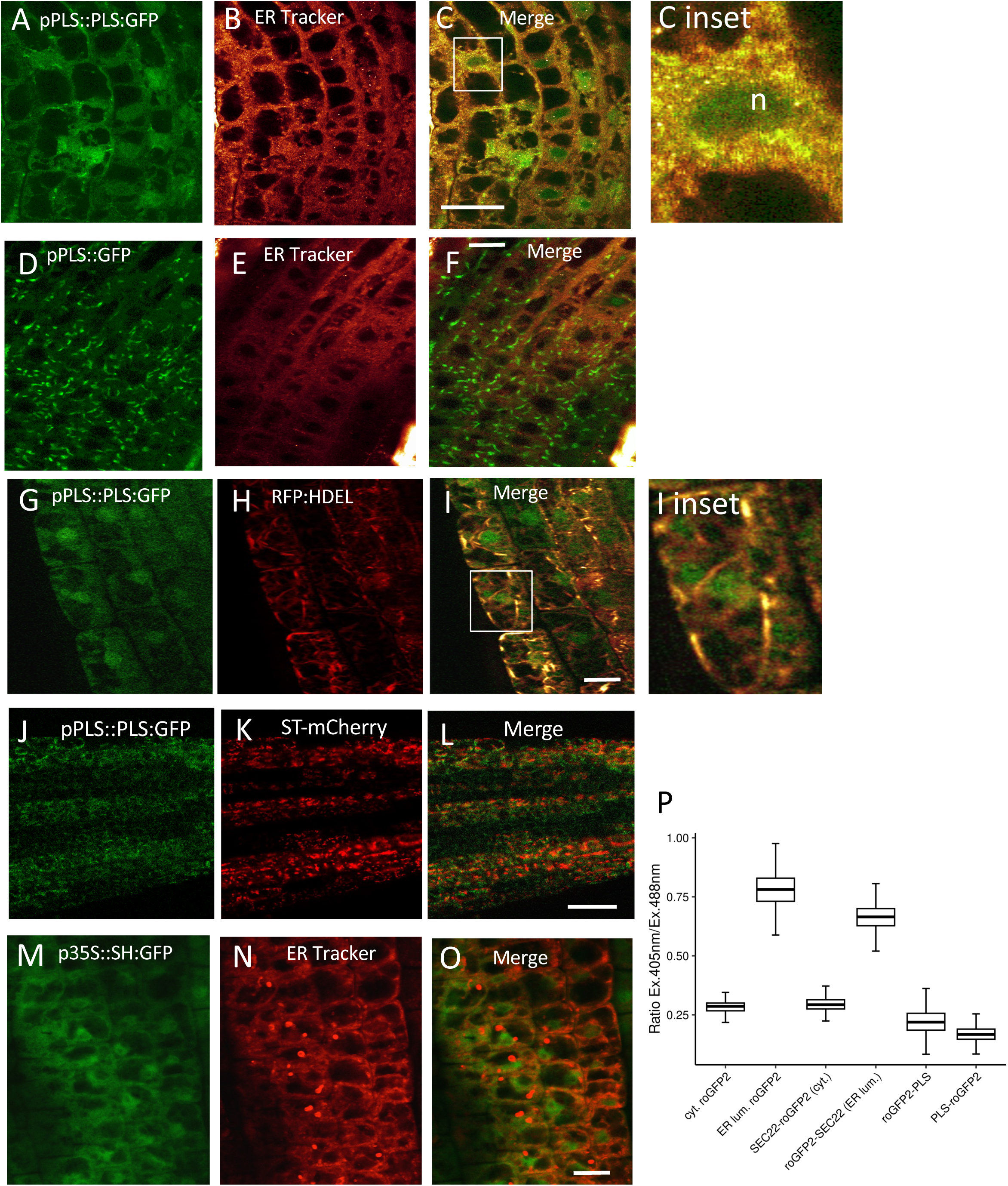
PLS localizes to the endoplasmic reticulum. (**A-O**) PLS::PLS:GFP fusion protein (**A, G, J**) colocalizes with endoplasmic reticulum markers ER Tracker (**B, C, C inset**) and RFP:HDEL (**H, I, I inset**), but free GFP does not (**D-F**, **M-O**). PLS:GFP staining is seen also in nuclei (n). PLS::PLS:GFP (**J**) does not co-localize with the *trans*-Golgi markers ST-mCherry SH:GFP (**K, L**). Scale bars = 25 μm (**C, l**), 10 μm (**F, I, O**). Root epidermal cells in the transition zone were imaged. (p) Ratiometric analysis of roGFP2 fusion constructs transiently expressed in *N. benthamiana*. Comparison of excitation ratios of PLS-roGFP2 and roGFP2-PLS with control constructs (free roGFP, SEC22 fusions) reveals that PLS localizes to the cytosolic side of the ER.

### PLS interacts with the ethylene receptor protein ETR1

We hypothesized that PLS plays a role in receptor function and investigated whether this involved direct interaction with the receptor complex. Preliminary experiments using yeast 2-hybrid analysis suggested that PLS interacts with ETR1 (Figure S5). Confirmation of the physical interaction between PLS and ETR1 in plants came from co-immunoprecipitation (Co-IP) analysis. *Agrobacterium* containing a plasmid encoding PLS linked to a C-terminal GFP and ETR1 with a C-terminal HA tag was infiltrated into *Nicotiana benthamiana* leaves for transient expression. After 3 d, interaction was confirmed by western blotting after Co-IP with either anti-GFP beads (showing PLS pulls down ETR1) or anti-HA beads (showing ETR1 pulls down PLS) (Figure 4A). GFP-only controls did not show binding with ETR1, demonstrating the interaction is dependent on the presence of the PLS peptide. The addition of 0.5 μM copper sulphate to the protein extract used for Co-IP experiments stabilized the PLS-ETR1 interaction. The presence of copper ions resulted in almost 3-fold more PLS:GFP detected upon pulldowns with ETR1-HA, or conversely of ETR1-HA pulled down with PLS-GFP, compared to the same assay in the presence of the metal chelator 2 mM EDTA (Figures 4A, B).

**Figure 4.**
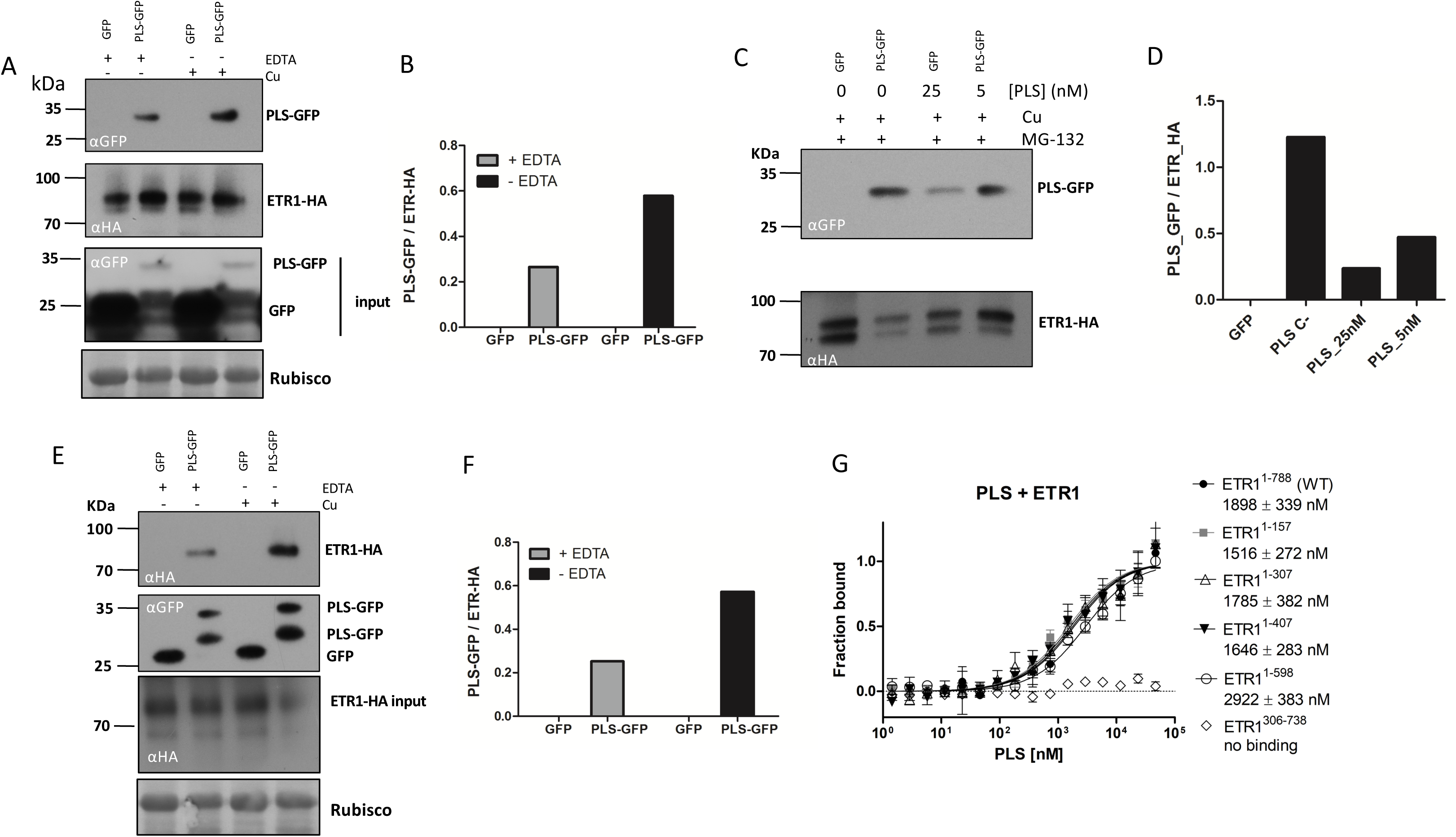
PLS interacts with the ethylene receptor ETR1. (A) Co-immunoprecipitation of PLS:GFP by ETR1:HA (upper panel) in leaves of *Nicotiana benthamiana*, in the presence and absence of 0.5 μM CuSO_4_ and EDTA (to remove Cu). Lower panels show presence of ETR1:HA in extracts using anti-HA antibody, and protein input blots using anti-GFP and anti-Rubisco. (**B**) Densitometric scan of immunoblot. (**C**) Competition assay showing a reduced binding between PLS:GFP and ETR1:HA in the presence of 0, 5 nM or 25 nM PLS peptide, in the presence of 0.5 μM CuSO_4_ and 50 μM MG-132, a proteasome inhibitor (upper panel). Lower panel shows ETR1:HA in extracts using anti-HA antibody. (**D**) Densitometric scan of immunoblot. (**E**) Co-immunoprecipitation of ETR1:HA (upper panel) in leaves of *N. benthamiana*, showing the effect of EDTA (to remove Cu) on interaction between ETR1:HA and PLS:GFP. Lower panels shows presence of ETR1:HA in extracts using anti-HA antibody, and protein input blots using anti-ETR1-HA and anti-Rubisco. (**F**) Densitometric scan of immunoblot. α-GFP, anti-GFP antibody; α-HA, anti-HA antibody. (**G**) Microscale thermophoresis binding curves of different ETR1 truncations with PLS. Binding of PLS was observed with full-length ETR1 and all C-terminal truncations but not with ETR1^306-738^ lacking the N-terminal transmembrane part of the receptor.

To investigate the specificity of PLS binding, synthetic full length PLS peptide PLS(FL) was introduced into infiltrated *N. benthamiana* leaves 30 min before tissue harvest in the presence of both copper to maximise PLS-ETR1 interaction and the proteasome inhibitor MG-132 to prevent protein degradation. The addition of 25 nM synthetic PLS caused a ca. 80% reduction in PLS-GFP binding to ETR1-HA (Figures 4C, D), suggesting that the synthetic PLS peptide competes for ETR1 binding, showing the specificity of PLS for ETR1. The anti-GFP beads bound two sizes of PLS-GFP protein (Figure 3E), both of which were larger than a GFP-only control, suggesting that the PLS peptide undergoes cleavage, a change in conformation, post-translational modification or incomplete reduction of Cys residues on some PLS. When using ETR1-HA to pull down PLS-GFP, only the larger peptide was present (Figures 4E, F), suggesting that ETR1 binds a longer version of the PLS peptide.

To pinpoint the interaction site at the receptor in more detail, *in vitro* binding studies were performed with purified receptor variants and PLS by microscale thermophoresis (MST; Figure 4G). Binding of PLS was observed only with receptor variants containing the N-terminal transmembrane domain (TMD). In contrast, no binding was detected with ETR1 lacking this domain (ETR1^306-738^). The TMD harbors the ethylene and copper binding region (Schott-Verdugo et al., 2019).

### PLS binds Cu(I)

Cysteine residues are common metal-ligand binding residues in low molecular weight copper-handling peptides, and predictions of PLS structure suggests a single a-helix plus unstructured region with two cysteines (CX_10_C arrangement where X is any amino acid), with some analogy to copper-metallochaperones such as Cox17 or other CX_9_C twin proteins (Robinson and Winge, 2010). In view of both structural considerations and the copper-dependency of ETR1 we determined whether the two cysteine residues (C6 and C17) in PLS play a functional role, by analysing both *pls* mutant complementation and in copper binding by PLS variants. A mutated *Arabidopsis* full-length peptide in which both cysteines were replaced with serines, PLS(FL C6S, C17S), was non-functional in hydroponic root feeding assays, failing to rescue the short primary root phenotype of the *pls* mutant (Figure 5A). Furthermore, as indicated above, the 9 amino acids sequence N2 and C-terminal sequences C1 and C2 from *Arabidopsis* PLS, which each contain only one cysteine residue, were unable to rescue the mutant (Figure 2C). These results indicate the requirement of the two cysteine residues for biological activity, measured as primary root growth. Interestingly, RNA-seq data showed that nineteen out of 287 genes associated with response to metal ion (GO:0010038) were also significantly down-regulated in the *pls* mutant (Table S4, Figure S2B, enrichment *FDR = 0.000057*; *20*).

**Figure 5.**
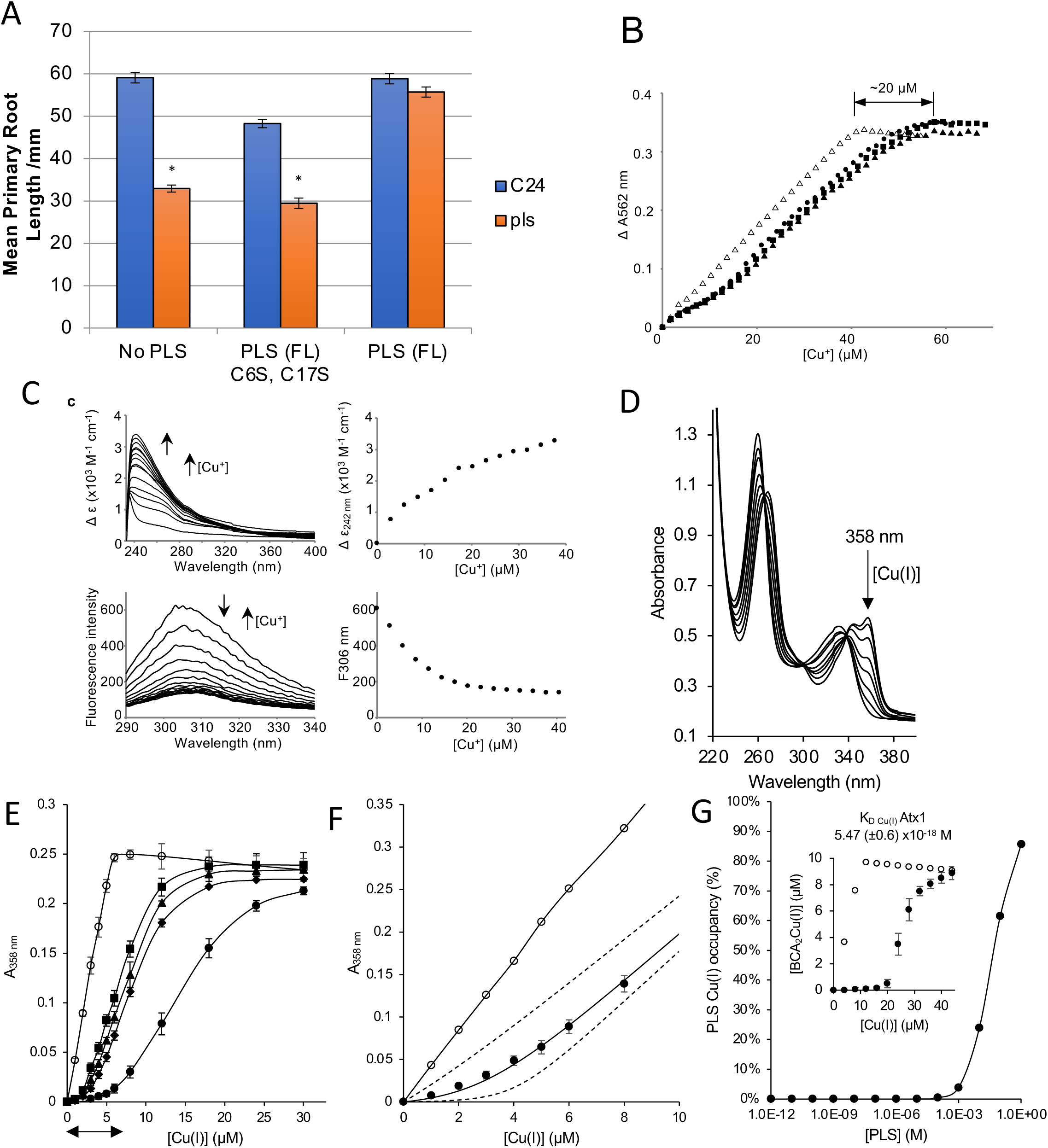
PLS binds copper and interacts with RAN1 and copper chaperones. (**A**) C6S and C17S in PLS are required for function. Seedlings were grown hydroponically for 10 d in the presence (100 nM) or absence of peptide (FL or C6S, C17S). Blue bars are C24, red bars are *pls*. *: *P* <0.05, *t*-test between C24 and *pls*, n = 25. Error bars show ± 1 standard error, n = 14-22. (**B**) BCA absorbance in the presence of synthetic PLS (44.8 μM, filled symbols) or an equivalent volume (to PLS) of DMSO (open symbols). Experiments with PLS used 88.3 μM (circles), 84 μM (triangles) and 88.4 μM (squares) BCA. The control experiment with DMSO used 85.3 μM BCA. (**C**) UV-vis apo subtracted difference spectra of synthetic PLS (19.8 μM) upon titration with CuCl (top left); binding isotherm of the feature at 242 nm (top right). Fluorescence spectra of PLS (19.8 μM) upon titration with CuCl (bottom left); binding isotherm of the feature at 306 nm (bottom right). (**D**) Absorption of BCA (17.3 µM) titrated with Cu(I) (representative spectrum, n = 2, full data in Fig. S6). (**E**) Binding isotherms (A_358 nm_) of BCA (10 µM) in the presence/absence (filled/empty symbols, respectively) of 14 µM MBP-PLS (circles) or MBP-PLS mutants (C6S, C17S and C6S/C17S: triangles, diamonds and squares respectively) titrated with Cu(I) (n = 3, ± SD). Arrow indicates ca. 7 µM withholding of Cu(I) from BCA. (**F**) Binding isotherms (A_358 nm_) of BCA (50 µM) in the presence/absence of 10 µM MBP-PLS (filled/empty symbols, respectively) titrated with Cu(I). Model (solid line) describes Cu(I)-binding as a 2:1 complex, with β*_2_* affinity of 3.79 (± 1.5) x10 M. Dotted lines simulate 10x weaker or tighter affinity (n = 3, ± SD). (**G**) Simulated Cu(I) occupancy (Supplementary Text) as a function of [PLS] using β*_2_* Cu(I) affinity of 3.79 x10^19^ M^-2^. Inset, *Arabidopsis* ATX1 (20 µM, filled circles) withholds one Cu(I) equivalent from 20 µM BCA (open circles, BCA-alone) (n = 3, ±SD), with *K_D_* Cu(I) 5.47 x10^-18^ M (Fig. S3).

To determine a possible role for PLS and the cysteine residues in binding Cu(I), synthesised PLS peptide was titrated with Cu(I) ions under strictly anaerobic conditions and monitored for copper-dependent spectral features (Figures 5B-D). Both UV-vis (putative metal-to-ligand charge-transfer) and fluorescence (putative tyrosine solvent access) spectra of PLS changed as a function of Cu(I) ion concentration, consistent with complex-formation (Figure 5C). We titrated PLS peptide against bicinchoninic acid (BCA; Figure 5B), which is a chromophore that binds Cu(I) (Xiao et al., 2011; Figures 5D and S6). The rationale was to determine whether PLS could compete Cu(I) from BCA, indicative of binding by PLS. PLS peptide was titrated with Cu(I) in the presence of 87 μM BCA. PLS peptide (44.8 μM) increased the concentration of Cu(I) required to saturate BCA increased by ca. 20 μM (Figure 5B), and the initial gradient is shallower than in the control of BCA and Cu(I) but lacking PLS. Thus PLS withholds Cu(I) from BCA, implying an affinity for Cu(I) (*K*_Cu_ PLS) tighter than *K*_Cu_ of BCA, and these data also provide support for a 2:1 stoichiometry of binding to the tightest site. Titration experiments also show that although synthetic PLS(FL) withholds Cu(I) from BCA (Figure S7), a mutant PLS (FL C6S, C17S) does not (Fig. S8), confirming a role for those cysteine residues in Cu(I) binding. As an additional and complementary strategy, to eliminate potential solubility problems of PLS in determining accurately the stoichiometry and affinity for Cu(I) (these studies require complete solubility for precise affinity quantification), we also tested a PLS fusion to maltose binding protein (MBP-PLS) which was found to retain solubility when titrated with Cu(I) ions under strictly anaerobic conditions. MBP-PLS (14 μM) was found to withhold ca. 7 μM Cu(I) from BCA, again indicative of a 2:1 PLS:Cu(I) stoichiometry, and tight binding is cysteine-dependent (Figure 5E). A β*_2_* affinity of 3.79 (±1.5) x10^19^ M^-2^ was determined by competition against an excess of BCA, and the fit significantly departs from simulations 10x tighter or weaker (Figures 5F, S9).

Metal binding in biology is challenging to predict because the formation of metal-protein complexes is a combined function of metal affinity for a given protein and metal availability, which would need to be known for Cu(I) in the *Arabidopsis* cytosol in the case of PLS (assuming the same orientation of binding residues as with the GFP tag; Figure 3P). Cu(I) occupancy of the cytosolic copper chaperone ANTIOXIDANT PROTEIN 1 (ATX1) tracks with fluctuations in available cytosolic Cu(I) such that its affinity approximates to the mid-point of the range of Cu(I) availabilities within this eukaryotic compartment (Yu et al., 2017; Morgan et al., 2019). *Arabidopsis* ATX1 was therefore expressed and purified to determine a 1:1 ATX1:Cu(I) stoichiometry and affinity *K*_D Cu(I)_ of 5.47 (±0.6) x10^-18^ M (Figure 5G inset, Figure S10). Figure 5F reveals that the cytosolic concentration of PLS would need to exceed (improbable) millimolar concentrations for Cu(I)-dependent homodimers to form at this cytosolic Cu(I) availability (mathematical models and equations shown in Supplementary text). It is thus unlikely that the Cu(I):PLS_2_ complex alone delivers Cu(I) to, or retrieves it from, the interacting cuproprotein ETR1. Cu(I)-dependent PLS-ETR1 heterodimeric complexes are the more likely functional species, and we conclude that PLS binding alters ETR1 conformation or ethylene availability to regulate receptor activity.

## DISCUSSION

We present evidence of a role for the PLS peptide as a new Cu(I)-binding peptide that, based on its physical interactions with the receptor, and its ethylene signalling function as determined by genetic and physiological experiments, acts as a regulatory component of the ethylene signalling pathway. Peptides can act as ligands for receptor kinases to regulate signalling pathways, such as in plant immunity (Campos et al., 2018), development (Willoughby and Nimchuk, 2021) or in response to abiotic stresses (Kim et al., 2021). We propose that the PLS peptide is a new component of the regulatory mechanism for ethylene receptor function linked to its copper-binding activity (Rodriguez et al., 1999). Loss of copper binding through mutation of the two cysteines in PLS ablates its biological activity (Figures 2B, C; Figure 5; Figure S8). The *pls* mutant phenotype has some similarities (enhanced ethylene responses) to strong *ran1* mutant alleles in which copper delivery to the receptor is compromised (Binder et al., 2010), and shares (statistically significant) overlap in DEGs with the *ctr1* mutant, whereby 11.9% of the *ctr1* DEGs are shared with *pls* (each compared to the respective wild types; Figure S11).

Many plant peptides involved in signalling are processed from longer pre-proteins (Olsson et al., 2019). In contrast, PLS is translated from an open reading frame to produce a functional 36 amino acids peptide; the peptide appears to be cleaved or otherwise modified in the cell (Figure 4E). It is possible that the three arginines (amino acids 10-12) may represent a cleavage site (Duckert et al., 2004; Capraro et al., 2015) which would produce the observed shorter GFP fusion protein (ca. 30 kD) detected in anti-GFP co-IP blots (Fig. 4E). The size of this PLS-GFP cleavage product suggests that this peptide fragment represents GFP (ca. 27 kDa) plus a 3 kDa fragment of the C-terminal region of PLS. Such cleavage would likely inactivate PLS function as we demonstrate that shorter N-and C-terminal fragments (each with only one cysteine residue) are not biologically active (Figure 2C). While PLS:GFP localizes to several subcellular compartments (though not the *trans*-Golgi), some localizes to the ER where the ethylene receptor is found (Figure 3). Furthermore PLS can interact directly with the receptor protein ETR1 (and specifically with the N-terminal copper-binding transmembrane domain of ETR1).

Previously there has been proposed a model for copper delivery into the cell via the COPT family of transporters, which are located at the root plasma membrane (Chen et al., 2022), to RAN1 via ATX1 and/or CCH (Li et al., 2017; Hoppen et al., 2019). Arabidopsis has three known copper chaperones, ATX1, CCH and a copper chaperone for superoxide dismutase CCS (Casareno et al., 1998; Chu et al., 2005; Puig et al., 2007). These are required for copper homeostasis, and both ATX1 and CCH have been shown to interact with RAN1 (Li et al., 2017), providing a link with ethylene signalling (Andrés-Colás et al., 2005; Puig et al., 2007). Chaperones are required for the transport of reactive copper to the correct compartment via these chaperones, avoiding cytotoxicity, following import into the cell; and into the xylem from the cytosol via HMA5, which is structurally similar to RAN1 (Williams and Mills, 2005; Burkhead et al., 2009). Loss of function mutants *cch* and *atx1* have no abnormal phenotype when grown under standard conditions, but the *atx1* and the *atx1 cch* double mutant, though not the *cch* single mutant, are hypersensitive to exogenous copper (Shin et al., 2012). ATX1 and CCH functions have likely diverged, a view supported by their differential transcriptional regulation by copper and different cell type specificities (Shin et al., 2012). The *ran1* mutant is however not copper-hypersensitive, while *hma5* is; and the *pls* mutant is, like *ran1*, not copper hypersensitive (Figure S12), indicative of a role that is distinct from that of ATX1 or CCH. Given its predicted low concentration in the cell and 2:1 stoichiometry with Cu(I) (Figures 5F and G, S9 and 10), we conclude PLS is unlikely to act as a cytosolic copper chaperone.

A lack of copper delivery in the strong *ran1-3* and *ran1-4* null alleles causes constitutive ethylene responses (Woeste and Kieber, 2000), producing a triple response phenotype similar to the *pls* mutant (Casson et al., 2002). These *ran1* mutants fail to bind ethylene due to the lack of copper at the ethylene binding site (Binder et al., 2010). The decrease in ethylene binding in the *ran1-3* and *ran1-4* mutants is not due to reduced levels of the ETR1 protein (Binder et al., 2010), and regulation of the ethylene receptor ETR1 is generally regarded to be independent from *ETR1* transcription or degradation (Hua et al., 1998). Weaker *ran1-1* and *ran1-2* alleles only show an ethylene response phenotype if they are grown in the presence of copper chelators (Binder et al., 2010). Therefore, defective RAN1 results in defective copper delivery to wild type receptors, generating non-functional receptors which cannot interact with CTR1, resulting in downstream ethylene responses.

Intriguingly, the *ran1-3* and *ran1-4* mutants with constitutive ethylene responses are distinct from the ethylene insensitive *etr1-1* mutant, which also cannot bind copper. Therefore, both the *etr1-1* mutant receptor and the strong *ran1* mutant alleles have reduced ethylene binding capability but the *etr1-1* mutation, which is gain-of-function, generates seedling phenotypes that are very different from seedlings carrying the ethylene hypersensitive wild type receptors that simply lack copper due to defective delivery (in *ran1-3* and *ran1-4).* A receptor carrying the *etr1-1* mutation is maintained in its ‘active’ state and constitutively inhibits downstream ethylene responses, whilst in the *ran1* mutants ETR1 remains in the ‘inactive’ state, promoting ethylene responses (Binder et al., 2010). Those authors therefore proposed that copper has more than one role in the regulation of ETR1 and ethylene signalling: it is crucial for binding ethylene molecules but it is also needed for the process of signal transduction by the receptor protein, by an unknown mechanism, to regulate downstream ethylene responses.

We propose that PLS represents a new component of this pathway, providing some tissue-specificity to ethylene receptor activity. Given the low probability that PLS would act as a cytosolic copper chaperone, it more likely regulates ethylene responses via its demonstrated Cu-dependent ETR1 binding. The RFP-HDEL and Rho-GFP co-localization experiments indicate the presence of PLS in both the cytosol and ER lumen (as well as other compartments; Figure 3).

Since the *pls* mutant is rescued by the supply of silver ions (Casson et al., 2002), and silver is likely delivered to the ethylene receptor via RAN1 (Binder et al., 2010), this argues against a mechanism in which PLS acts downstream of RAN1 to metalate ETR1 in the ER lumen. This would also be consistent with the data suggesting PLS is located predominantly at the cytosolic side of the ER; but it binds to the transmembrane domain of ETR1. It is also unlikely that PLS removes Cu(I) from the receptor as this would inactivate the receptor and lead to enhanced ethylene signalling in the presence of PLS, which is the opposite of what would be expected from the *pls* mutant phenotype. We therefore favour a model (Fig. 6) in which PLS binds ETR1 as part of the receptor biogenesis mechanism, with the PLS-ETR1 physical interaction enhanced in the presence of copper (Fig. 4), to regulate receptor conformation to generate the active form, in which ethylene responses are repressed. This may be through regulation of Cu(I) homeostasis at the receptor, or via other structural interactions - for example, PLS could physically block ethylene binding at the Cu(I) pocket. This question can form the basis of future investigations. If PLS is required for Cu(I) delivery, it may act redundantly with RAN1, accounting for the relatively mild ethylene phenotype of the *pls* mutant (compared with e.g. *ctr1*).

**Figure 6.**
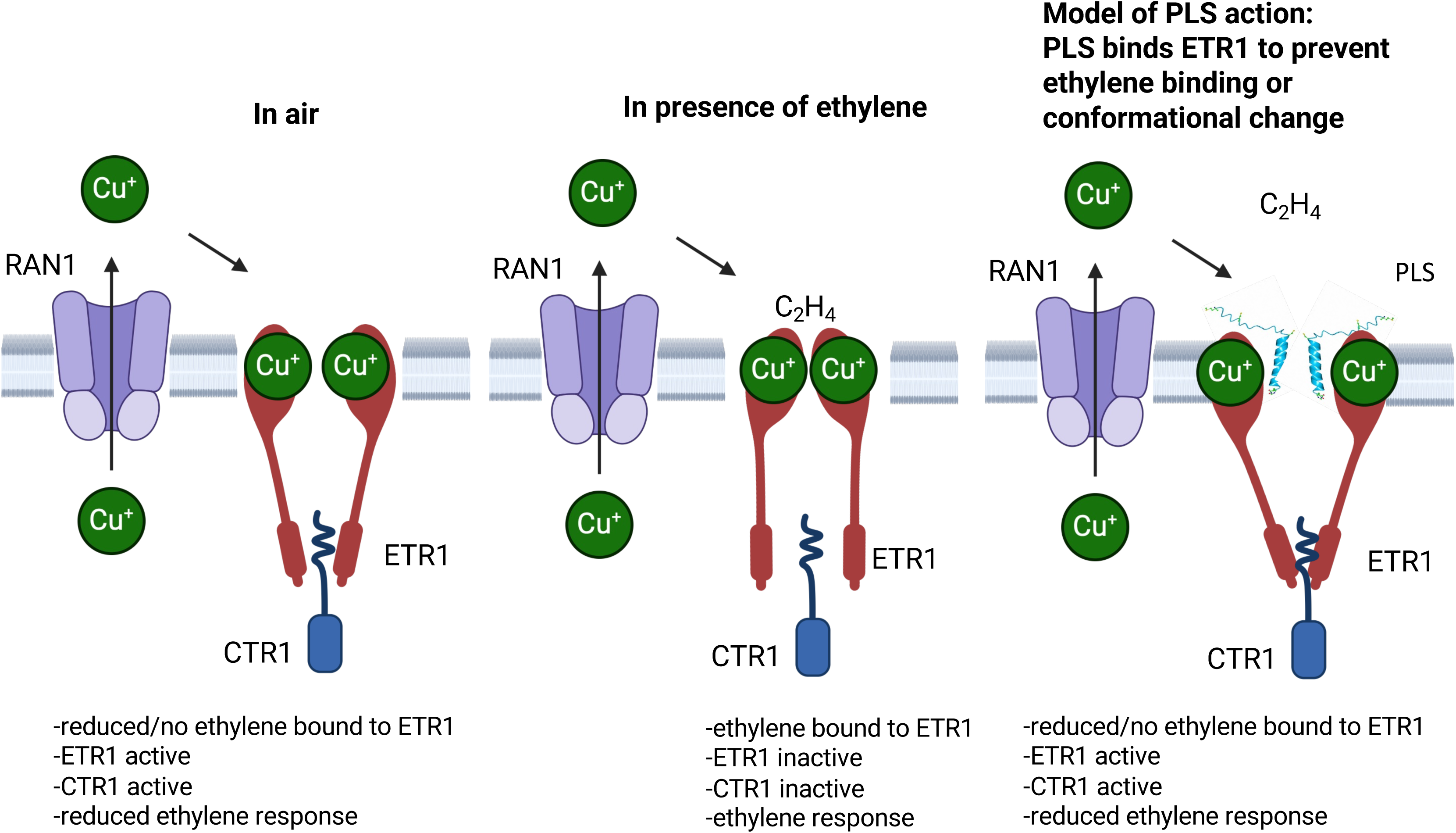
Model of the role of PLS in ethylene signalling. In this model, we assume that RAN1 and ETR1 are in the same membranous compartment (likely the ER) at least transiently. RAN1 is believed to pump Cu(I) into the ER lumen, and then to ETR1. In air (absence of ethylene; left panel), the receptor complex is active and activates CTR1, which is a negative regulator of ethylene responses. In the presence of ethylene (centre panel), the receptor is in an inactive state, cannot activate CTR1, resulting in ethylene responses. In the Model for PLS function, PLS binds to ETR1 (with binding potentially enhanced by the Cu(I) in the ETR1 transmembrane domain) and affects the receptor conformation, potentially blocking ethylene binding to ETR1 or inducing some other conformational change, leading to active ETR1, active CTR1 and reduced ethylene signalling. This model is consistent with the observed *pls* loss of function mutant showing enhanced ethylene responses.

Given that *PLS* is expressed in specific tissues, notably root tip and leaf vascular tissue (Casson et al., 2002), it seems likely that PLS adds a level of regulation of receptor function in specific tissues, with a focus in this paper on the root (though developmental defects in the vascular patterning of the leaf are also observed in the *pls* mutant; Casson et al., 2002). We propose that PLS acts to dampen ethylene responses at sites of high auxin. *PLS* transcription is strongly up-regulated by auxin, and its expression profile is similar to that of the auxin reporter DR5 (Sabatini et al., 1999; Casson et al., 2002). The root tip, where *PLS* is most strongly expressed, is a site of high auxin concentration, whereby an auxin maximum is formed around the stem cell niche to regulate both stem cell identity and function (Sabatini et al., 1999). The vascular tissues are also a site of high auxin (Bishopp et al., 2011). Auxin can induce ethylene biosynthesis by upregulation of the ethylene biosynthetic enzyme ACC synthase (Abel et al., 1995), and high ethylene concentrations can lead to both reduced meristem function and aberrant division of the quiescent centre and cells within the stem cell niche and columella (Ortega-Martinez et al., 2007). Therefore ethylene biosynthesis and signalling in the root tip must be tightly regulated to allow growth control. *PLS* expression is transcriptionally activated by auxin and suppressed by ethylene. We have shown in previous work that there is crosstalk between PLS, ethylene and other signalling pathways, notably auxin, cytokinin and ABA in the root (e.g. Liu et al., 2010, 2013; Moore et al., 2015), and the distinctiveness of *pls* compared to other ethylene signaling mutants may be linked to its restricted expression pattern. This network provides a mechanism to suppress growth-inhibitory ethylene responses in the high auxin environment of the root tip, which is dependent on the tissue-specific expression of *PLS* (Liu et al., 2010) (Figure S13) and is mediated through its interaction with the ethylene receptor.

Future structural studies should reveal more about the PLS-ETR1-RAN1-chaperone interaction and diverse roles of copper ions in ethylene signal transduction, which represents a new paradigm for the regulation of signaling protein function by metals.

## METHODS

### Plant Material

Seedlings of *Arabidopsis thaliana* ecotype either C24 or Col-0, or *pls* mutants or transgenic *PLS* overexpressers, all from lab stocks, were grown on solid sterile half-strength MS10 medium on 2.5% Phytagel (Sigma-Aldrich) in 90 mm Petri dishes (Sarstedt, Leicester, UK) at 21°C, under a 16 hour photoperiod as described (Casson et al., 2002; Chilley et al., 2006). *Nicotiana benthamiana* plants (lab stocks) were grown in a controlled environment (21°C, 16 h photoperiod) for transient expression studies in leaf tissue. For hydroponic feeding studies, *Arabidopsis* seedlings were cultured in liquid medium (1 ml/well) in sterile 24-well plates (Sarstedt), essentially as described (Matsuzaki et al., 2010). For hormone and peptide assays, one seedling was grown in each well, at 21°C, under a 16 h photoperiod. For peptide feeding experiments, purified freeze-dried peptide was dissolved in DMSO to create a 500 μM stock solution. Peptide stock solution was added to liquid ½ MS10 plant media containing 0.1% DMSO to make a final peptide concentration of 50 or 100 nM (or 10, 25, 50 and 100 nM for dose-dependent assays). For copper treatments, 1 mM CuSO_4_ solution was filter-sterilized and added to autoclaved liquid ½ MS10 plant media to create final CuSO_4_ concentrations of 0, 5, 10, 15, 20, 25, 30, 35, 40, 45 and 50 μM. The copper chelator bathocuproine disulphonic acid (BCS) was added to liquid medium to produce final concentrations of 0, 10, 50, 100, 250 and 500 μM. Seedlings were scanned to create a digital image and root lengths of seedlings were measured using ImageJ. Statistical analysis was performed using the Real Statistics Resource Pack software (Release 3.8, www.real-statistics.com) in Excel (Microsoft).

### Peptide Synthesis

Peptides were either obtained from Cambridge Research Biochemicals (Billingham, UK) or synthesized in the laboratory by Fmoc solid phase peptide synthesis (SPPS) on a CEM Liberty1 single-channel peptide synthesiser equipped with a Discover microwave unit. Reactions were performed in a 30 ml PTFE reaction vessel with microwave heating FW: EpiLife Media and agitation by bubbling nitrogen. Peptide synthesis was carried out using 2-chlorotrityl chloride resin (0.122 mmol g^-1^, Novabiochem) using Fmoc-protected amino acids. The first amino acid residue at the C-terminus (histidine) was coupled manually by mixing 76 mg (1 eq.) Fmoc-His(Trt)-OH, 0.09 ml (4 eq.) *N,N*-diisopropylethylamine (DIPEA), 1 ml dichloromethane (DCM) and 1 ml dimethylformamide (DMF) until the amino acid powder had dissolved. The mixture was added to 0.1 mmol resin and stirred gently for 120 min at room temperature. Resin was washed with 3x DCM/MeOH/DIPEA (17:2:1), 3x DCM, 2x DMF and 2x DCM. Amino acid coupling reactions were performed using Fmoc-protected amino acids present in a 5-fold excess (2 M concentration), HOBt (0.5 M HOBt in DMF, used at the activator position) and DIC (0.8 M in DMSO, used at the activator base position). For double and triple couplings the reaction vessel was drained after each coupling cycle and fresh reagents were added. Before each coupling, a room temperature preactivation period of 1 to 2 h was used. Microwave-assisted couplings were performed for 10 min at 75°C at 25W power. Cys and His residues were coupled at low temperature (10 min at room temperature followed by 10 min at 50°C, 25W). Arg residues were double coupled, firstly by 45 minutes at room temperature plus 5 minutes at 75°C (25W), and second by the standard microwave conditions above. Fmoc group removal was carried out by two piperidine solution treatments (20% piperidine in DMF) in succession: 5 minutes then 10 minutes. Peptide cleavage from resin was carried out in 3 ml 95% TFA in dH_2_O/TIPS (2.85 ml TFA, 0.15 ml dH_2_O, 0.15 ml triisopropylsilane). Peptide was dissolved in water with a small volume of MeCN and lyophilized to produce a powder using a Christ ALPHA 1-2 LD_plus_ freeze dryer.

### Preparative High-Performance Liquid Chromatography (HPLC)

Peptide products were analysed and purified by HPLC at 280 nm. 25-50 mg of freeze-dried peptide sample was dissolved in 1 ml 1:1 H_2_O:MeCN and injected onto a Speck and Burke Analytical C18 Column (5.0μm, 10.0 x 250 mm) attached to a PerkinElmer (Massachusetts, USA) Series 200 LC Pump and 785A UV/Vis Detector. Separation was achieved by gradient elution of 10-80% solvent B (solvent A = 0.08% TFA in water; solvent B = 0.08% TFA in ACN) over 60 min, followed by 80-100% B over 10 min, with a flow rate of 2 ml/min. Selected peptide fractions were lyophilized and a mass assigned using MALDI-TOF MS. Peptide sequences were identified using MALDI-TOF MS, using an Autoflex II ToF/ToF mass spectrometer (Bruker Daltonik GmBH, Germany) equipped with a 337 nm nitrogen laser. MS data was processed using FlexAnalysis 2.0 (Bruker Daltonik GmBH).

### Imaging

CSLM images for propidium iodide and GFP were obtained by a Leica SP5 TCS confocal microscope as previously described (Rowe et al., 2016). *pPLS::PLS:GFP, pPLS::GFP* and *p35S::GFP* seedlings were grown for 7 d on Phytagel ½ MS10 medium before ca. 25 mm of the root tip was removed and mounted in dH_2_O prior to imaging. The ER marker *p35S::RFP:HDEL* (*24*) (provided by Dr. Pengwei Wang, Durham University), and the trans-Golgi apparatus marker *pFGC-ST:mCherry* (from Nottingham Arabidopsis Stock Centre, www.arabidopsis.info) were introduced into *pPLS::PLS:GFP* plants by the floral dip method of transformation (Clough and Bent, 1998) using *A. tumefaciens* GV3 101. ER was also localized using ER Tracker™ Red (Thermofisher). Seven day-old seedlings were stained for 30 min in the dark in liquid ½ MS10 media containing 1 μM ER Tracker™ Red.

### RNA Isolation, RNA Sequencing, RT-qPCR

RNA was extracted from 7 day-old seedlings grown on half strength MS10 medium essentially as described (Thompson et al., 2023). The Illumina HiSeq 2500 System was used for RNA sequencing of three biological replicate samples, with libraries prepared using the Illumina TruSeq Stranded Total RNA with Ribo-Zero Plant Sample Preparation kit (RS-122-2401), essentially as described (Thompson et al., 2023). Library quality control was carried out again using a TapeStation with D1000 ScreenTape (catalog number 5067-5582). RNA-seq data were aligned to the TAIR10 (EnsemblePlants, release 58) genome sequence with corresponding gtf file using STAR (v 2.7.11a; Dobin et al., 2013) to obtain read count per gene. Read count per gene was analysed using DESeq2 v 1.40.2 (Love et al., 2014) to get p values, adjusted p values, and log2 fold changes. Differentially expressed genes (DEGs) were identified with the criteria of adjusted p value < 0.05. Gene ontology analysis was performed using AgriGO (Tian et al., 2017) singular enrichment analysis. Gene expression heatmaps were generated from log2 normalized counts using pheatmap (v1.0.12, https://cran.r-project.org/web/packages/pheatmap/index.html) with row scaling. RNA-seq data were deposited in GEO (https://www.ncbi.nlm.nih.gov/geo/) with accession number GSE256166 (Mudge et al., 2024).

For RT-qPCR, RNA was extracted from 7 day-old seedlings (3 biological replicates, 20 mg of tissue) as described (Thompson et al., 2023). Samples were checked for the presence of genomic DNA by PCR with *ACTIN2* primers ACT2 forward and reverse. Primer sequences were determined using Primer-BLAST (https://www.ncbi.nlm.nih.gov/tools/primer-blast/). Primers are listed in Table S7.

### Protein-protein interaction studies

#### Yeast 2-hybrid

The GAL4 two-hybrid phagemid vector system was used to detect protein-protein interactions *in vivo* in yeast, using the reporter genes β-galactosidase *(lacZ)* and histidine (*HIS3*) in the YRG-2 yeast strain, essentially as described (Zhong et al., 2008). DNA that encodes the target (ETR1) and bait (PLS) was inserted into the pAD-GAL4-2.1 A and pBD-GAL4 Cam phagemid vectors respectively and expressed as hybrid protein. The hybrid proteins were then assayed for protein-protein interaction.

DNA encoding the target and bait proteins were prepared by PCR amplification using primer designed the specifically for the DNA encoding the target (ETR1) and bait (PLS). Each set of primer contained specific endonucleases on the ends of primer corresponding to the endonucleases in the MCS of pAD-GAL4-2.1 A and pBD-GAL4 Cam phagemid vectors. The DNA construct of the target (ETR1) and bait (PLS) with specific restriction sites on the ends was then transformed into the TOPO 2.1 vector to check the sequence of the amplified DNA by sequencing with M13 forward (CTG GCC GTC GTT TTA C) and M13 reverse (CAG GAA ACA GCT ATG AC). The two vectors, pAD-GAL4-2.1 and pBD-GAL4 Cam were digested using specific restriction endonucleases and dephosphorylated prior to ligating the insert DNA. The DNA encoding the target (ETR1) and bait (PLS) was then ligated into the same reading frame as the GAL4 AD of the pAD-GAL4-2.1 phagemid vector and the GAL4 BD of the pBD-GAL4 Cam phagemid vector.

The following primers were used for PCR amplification:

ETR1:

Forward primer GAA TCC ATG GAA GTC TGC AAT TGT A (Eco RI on 5’ end) Reverse primer GTC GAC TTA CAT GCC CTC GTA CA (Sal I on 5’end)

PLS:

Forward primer CTG GAG ATG AAA CCC AGA CTT TGT (Xho I on 5’ end) Reverse primer GTC GAC ATG GAT TTT AAA AAG TTT (Sal I on 5’ end)

The pGAL4 control plasmid was used alone to verify that induction of the *lacZ* and *HIS3* genes has occurred and that the gene products are detectable in the assay used. The pLamin C control plasmid was used in pairwise combination with the pAD-WT control plasmid or with the pAD-MUT control plasmid to verify that the *lacZ* and *HIS3* genes are not induced as the protein expressed by each of these pairs do not interact *in vivo*.

Control plasmids were transformed into the YRG-2 strain prior to the initial transformation of the bait and the target plasmids and used separately or in pairwise combination in the transformation of the YRG-2 yeast strain. Yeast competent cells were cotransformed with the bait and target plasmids by sequential transformation.

### Gene cloning for co-immunoprecipitation

To investigate the interaction between the PLS peptide and the ethylene receptor ETR1, two DNA constructs were created by Gateway cloning. The 105-nucleotide *PLS* gene (without the stop codon) was inserted into the pEarlyGate103 (pEG103) destination vector, containing the *p35S* promoter and a C-terminal GFP tag, producing a vector containing the *p35S::PLS:GFP* DNA. The ETR1 cDNA was inserted into the pEarlyGate301 (pEG301) vector to create a *p35S::ETR1:HA* construct, producing an ETR1 protein with a C-terminal HA tag.

### Infiltration into Nicotiana benthamiana

The transient expression of constructs in *Nicotiana benthamiana* (tobacco) leaves was as previously described (Voinnet et al., 2007). Experiments were replicated up to 5 times. Competent *Agrobacterium tumefaciens* GV3101 cells were transformed with the desired plasmid containing the gene of interest and each injected with a syringe. The plants were approximately 7 to 10 weeks old; the chosen leaves were healthy and of length 3-6 cm, and 3 to 4 leaves were infiltrated with each construct.

### Protein extraction and PLS/ETR1 co-immunoprecipitation

Total protein was extracted from the infiltrated leaves of *N. benthamiana* plants 3 d after infiltration for co-immunoprecipitation experiments to investigate the interaction between PLS and ETR1 proteins, essentially as described previously (Srivastava et al., 2020). For competition assays, 5 nM or 25 nM full-length PLS peptide was also infiltrated in the presence of 50 μM MG-132 (a proteasome inhibitor) 30 min prior to tissue freezing. ChromoTek (Planegg, Germany) anti-GFP beads were used to immunoprecipitate the PLS-GFP protein, and Sigma-Aldrich (St. Louis, USA) anti-HA beads for the HA-tagged ETR1.

SDS-PAGE was used to separate protein fragments. The complexed proteins from the pull-down assay were analysed on 10-12% acrylamide gels. Membranes were incubated with primary antibody for 2.5 h (GFP, Abcam, Cambridge, UK: rabbit, 1:10000; HA (Roche), rat, 1:3000; Rubisco large subunit (Agrisera), rabbit, 1:10000). Excess primary antibody was then removed by washing three times in 2x TBST (150 mM NaCl, 10 mM Tris, 0.1% (v/v) Tween 20, pH 7.4) for 2 m, 5 min and 10 min, and subsequently incubated for 1 h with the ECL peroxidase-labelled anti-rabbit or anti-rat IgG secondary antibody, diluted 1:20000 in TBST. Excess secondary antibody was removed again by washing three times in 1x TBST, as with the primary antibody. In order to visualize the probed blot, the membrane was incubated with ECL Western Blotting Detection Reagent immediately prior to imaging. The horseradish peroxidase conjugated to the secondary antibody was detected by using X-ray film.

### Estimation of synthetic PLS concentration

Freeze-dried synthetic PLS peptide (Cambridge Research Biochemicals, Billingham) was dissolved in DMSO. An aliquot was added to aqueous buffer (10 mM HEPES pH7, 20 mM NaCl, 80 mM KCl) and absorbance at 280 nm was recorded. Concentration was estimated from the absorbance and the ProtParam estimated extinction coefficient of 2,980 M^-1^ cm^-1^. Concurrent with this a sample was submitted for quantitative amino acid analysis (Abingdon Health Laboratory Services). From this analysis a conversion factor of 2.27 was generated, which was applied to concentrations determined by A280 nm.

### ATX1 purification

*E. coli* BL21(DE3) containing pETatx1 was used to overexpress the wild type *ATX1* gene from *Arabidopsis thaliana* (optimised for expression in *E. coli*, Novoprolabs). Harvested cells were collected and frozen at-20 °C overnight then defrosted, resuspended in 20 mM HEPES (pH 7.0), 10 mM EDTA, 100 mM NaCl, 10 mM DTT. Cells were sonicated (Bandelin Sonoplus), supernatant separated by size exclusion chromatography (GE Healthcare, HiLoad 26.600 Superdex 75 pg) metal-free buffer lacking EDTA. Fractions containing ATX1 were incubated overnight and pooled before transfer into an anaerobic chamber (Belle Technology) via desalting column where reductant was removed. ATX1 was quantified by combination of Bradford assay and Ellman’s reagent to ensure the fully reduced state of the protein. Samples were also analysed for metal content by ICP-MS to ensure > 95% apo-ATX1.

### MBP-PLS/mutant purification

A fusion of PLS to MBP was created using the NEBExpress MBP Fusion and Purification System. Two complementary oligonucleotide primers encoding PLS (optimised for expression in *E. coli*) were annealed and inserted into the pMal-c5x plasmid at XmnI and SalI insertion sites.

The three mutants MBP-PLS(C6S), MBP-PLS(C17S) and MBP-PLS(C6S/C17S) were created by site-directed mutagenesis (QuikChange II, Agilent).

*E. coli* NEB Express containing the pMal plasmid with the correct MBP-PLS mutant was used to overexpress each protein. Harvested cells were resuspended in 20 mM Tris-HCl (pH 7.4), 200 mM NaCl, 1 mM EDTA and frozen at-20 °C overnight. Cells were defrosted in cold H_2_O, sonicated, purified by ammonium sulphate precipitation (where MBP-PLS precipitates >60% saturation), separated on MBP-trap (GE Healthcare) and eluted using buffer containing 10 mM maltose. MBP-PLS-containing fractions were pooled and concentrated using centrifugal concentrator (Corning, Spin-X UF 30 KDa) and buffer exchanged by desalting column into metal-free 20 mM HEPES (pH 7.0), 50 mM NaCl buffer in an anaerobic chamber (Belle Technology). Mutants containing thiols were quantified Ellman’s assay and MBP-PLS(C6S/C17S), which lacks all thiols, was quantified by Bradford assay alone. Samples were also analysed for metal content by ICP-MS to ensure >95% apo-protein.

### Anaerobic spectroscopic analysis of Cu(I) complexes

All Cu(I) titration experiments were carried out in an anaerobic chamber (Belle Technology) using metal-free CHELEX-treated, degassed buffers. For experiments titrating Cu(I), aqueous CuSO_4_ stock was quantified in advance by ICP-MS and diluted to working concentrations. The reductant NH_2_OH was included at final concentration of 1 mM to retain Cu(I) in its reduced state. Proteins were diluted in buffer to the final concentration specified in each titration in air-tight quartz cuvettes (Helma), and after addition of probe to the concentration specified, titrated with CuSO_4_. After each addition, solutions were thoroughly mixed and absorbance spectra recorded using a Lambda 35 UV/Vis spectrophotometer (Perkin Elmer). Titration isotherm data was fitted using simulated affinity curves using DynaFit (Kuzmič, 2009).

### Interaction studies of PLS with copper transporter ETR1 by Microscale Thermophoresis

Fluorescently labelled ETR1 truncation mutants were added to a dilution series of PLS in 50 mM HEPES, 150 mM NaCl, 0.015 % (wv) FosCholine 16 (pH 7.6) or 50 mM Tris, 300 mM NaCl, 0.015 % (w/v) FosCholine 16 (pH 7.6). Dissociation constants were calculated using GraphPad Prism 5.

### Determination of dissociation constants for the PLS-ETR1 interaction

Full-length ETR1 and truncation mutants were purified and labelled as described in Milić e., 2018). 94 µM PLS were diluted serially in 50 mM Tris and 300 mM NaCl (pH 7.6). Fluorescently labelled receptor was added at a final concentration of 50 nM. Thermophoretic behaviour was measured in premium capillaries at 50 % LED and 50 % MST power. In case of a binding event, data were fitted using GraphPad Prism 5.

### Ratiometric analysis of roGFP2 fusion proteins

N-and C-terminal roGFP2 fusion proteins of PLS were generated by Gateway cloning. Infiltration and transient expression of roGFP2 fusions and control proteins were carried out as described (Brach et al., 2009). Image acquisition and data analysis were carried out as described by Hoppen et al. (2019). A minimum of 10 leaf optical sections were imaged and used for ratiometric analysis of the redox sensitive excitation properties of roGFP2.

## SUPPLEMENTAL INFORMATION

All study data are included in the article and/or Supplemental Information are available at *Molecular Plant Online*. RNA-seq and root assay data are available at GEO accession GSE256166: https://eur01.safelinks.protection.outlook.com/?url= https%3A%2F%2Fwww.ncbi.nlm.nih.gov%2Fgeo%2Fquery%2Facc.cgi%3Facc%3DGSE256166&data=05%7C02%7Ckeith.lindsey%40durham.ac.uk%7C9637f7022a2240cfb10608dc324244cd%7C7250d88b4b684529be44d59a2d8a6f94%7C0%7C0%7C638440507397042151%7CUnknown%7CTWFpbGZsb3d8eyJWIjoiMC4wLjAwMDAiLCJQIjoiV2luMzIiLCJBTiI6Ik1haWwiLCJXVCI6Mn0%3D%7C0%7C%7C%7C&sdata=U4JHCoviYbdEbF76Tws8Ridq7W%2B5Hq2qBVN%2BFjIysgw%3D&reserved=0

## AUTHOR CONTRIBUTIONS

KL and NJR initiated the project. KL, JFT, NJR, AS and GG designed and supervised aspects of the project. AJM, SM, WM, BO-P, WS, WW, CT, DR, FMH, CH and BU carried out the experimental work and prepared the Figures. KL, NJR and GG drafted the early version of the manuscript, and all authors reviewed and edited the manuscript.

## ACKNOWLEDGEMENTS

The authors declare no competing interests.

