## Supplementary text and Figures for "POLARIS is a copper-binding peptide required for ethylene signalling control in *Arabidopsis*" (from another version of this preprint: version 5, 2024-12-18)

**Derivation of equations used to describe the metalation of a protein (P) that forms a 2:1 complex ($MP_2$) with a metal (M).**

Cu(I) occupancy of the cytosolic copper chaperone ATX1 is tuned to track with fluctuations in available cytosolic Cu(I) such that its affinity provides a first approximation of Cu(I) availability within this compartment. Based on the determined Cu(I) stoichiometry of *Arabidopsis* ATX1 (Fig. 4C) and subsequent affinity (Fig. S10), Cu(I) occupancy of a 1:2 Cu(I):$PLS_2$ complex has been calculated as a function of total cytosolic [PLS].

The following equilibrium describes the formation of a 1:2 complex of metal (M) and protein (P)

$$M + 2P \leftrightharpoons MP_2$$

The equilibrium constant for the association of the reaction is

$$\beta_2 = \frac{[MP_2]}{[M][P]^2} \qquad (1)$$

The fractional occupancy of the protein with the metal M can be defined as:

$$Fractional\ occupancy\ (\%) = 100 \text{ x } \frac{2[MP_2]}{[P]_{tot}} \qquad (2)$$

The following is the mass balance equation for the protein

$$[P]_{tot} = [P] + 2[MP_2] \qquad (3)$$

Equations *(2)* and *(3)* can be combined to give

$$\frac{2[MP_2]}{[P]_{tot}} = \frac{2[MP_2]}{[P]+2[MP_2]} \qquad (4)$$

Substituting *(1)* into *(4)* gives

$$\frac{2[MP_2]}{[P]_{tot}} = \frac{2\beta_2[M][P]^2}{[P]+2\beta_2[M][P]^2} \qquad (5)$$

Simplifying leaves the fractional occupancy in terms of metal affinities ($\beta_2$), available metal concentration ([M]), and concentration of *apo-* (uncomplexed) protein ([P]).

$$\frac{2[MP_2]}{[P]_{tot}} = \frac{2\beta_2[M]}{[P]^{-1}+2\beta_2[M]} \qquad (6)$$

The *apo*-protein concentration is itself a function of metal affinities, total protein concentration and available metal concentration. Substituting *(1)* into *(3)* and rearranging gives

$$[P]_{tot} = [P] + 2\beta_2[M][P]^2$$

$$So: 2\beta_2[M][P]^2 + [P] - [P]_{tot} = 0 \qquad (7)$$

Can solve this equation for [P] using the quadratic formula:

$$[P] = \frac{-b \pm \sqrt{b^2 - 4ac}}{2a}$$

*(8)*

Where: $a = 2\beta_2[M],\ b = 1\ and\ c = -[P]_{tot}$

The fractional occupancy of the protein with metal can therefore be expressed as a function of metal affinity, metal availability and total protein concentration (*ie* measurable parameters) by combining equations (*6*) and (*8*)

$$\frac{2[MP_2]}{[P]_{tot}} = \frac{2\beta_2[M]}{\frac{-b \pm \sqrt{b^2 - 4ac}}{2a}^{-1} + 2\beta_2[M]} \qquad (9)$$

Where: $a = 2\beta_2[M],\ b = 1\ and\ c = -[P]_{tot}$

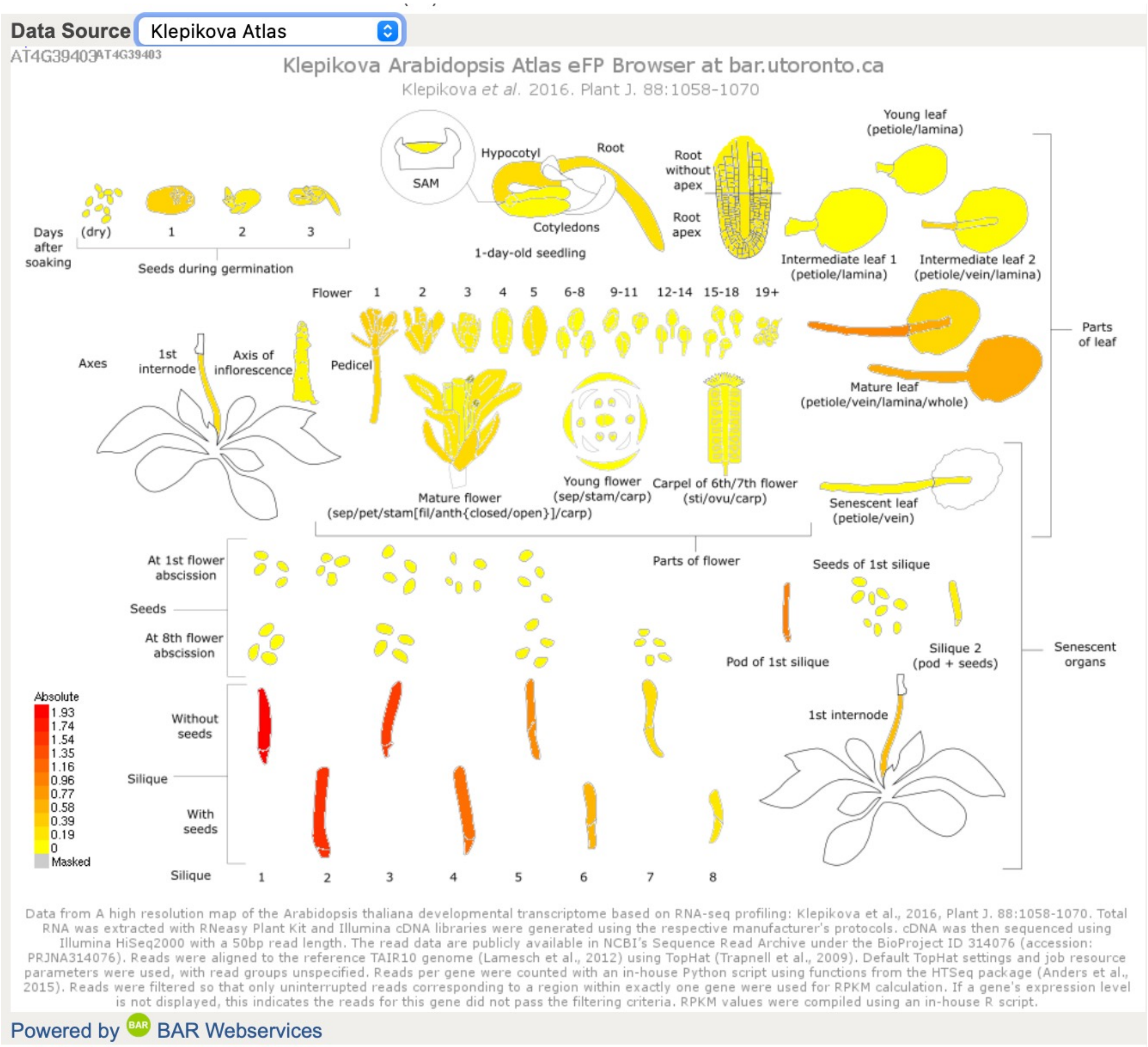


**Figure S1. *POLARIS* (AT4G39403) expression.**
A high resolution map of the Arabidopsis thaliana developmental transcriptome based on RNA-seq profiling: Klepikova et al., 2016, Plant J. 88:1058-1070.

From https://www.arabidopsis.org/servlets/TairObject?id=1001029664&type=locus

A

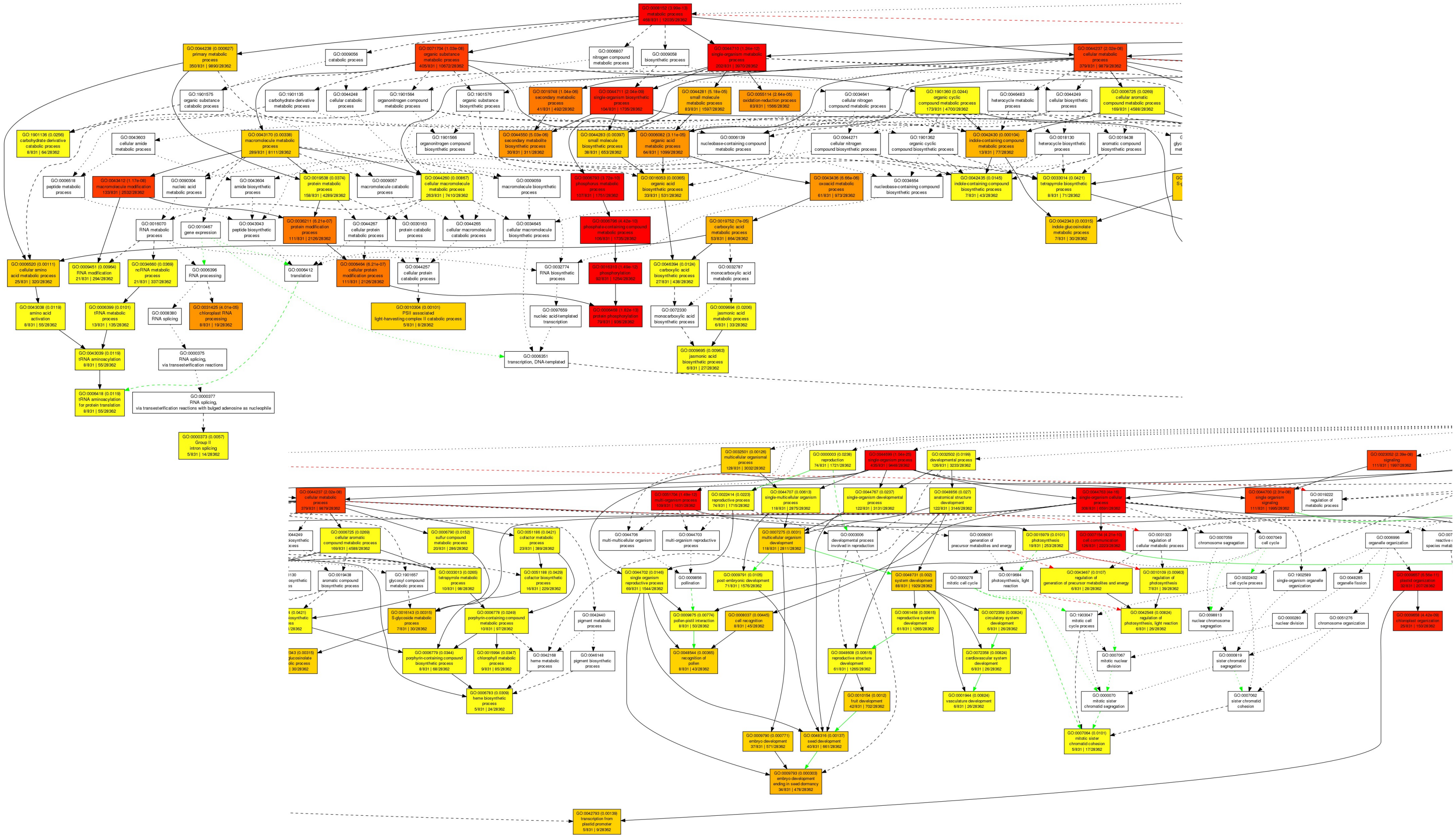

A contd.

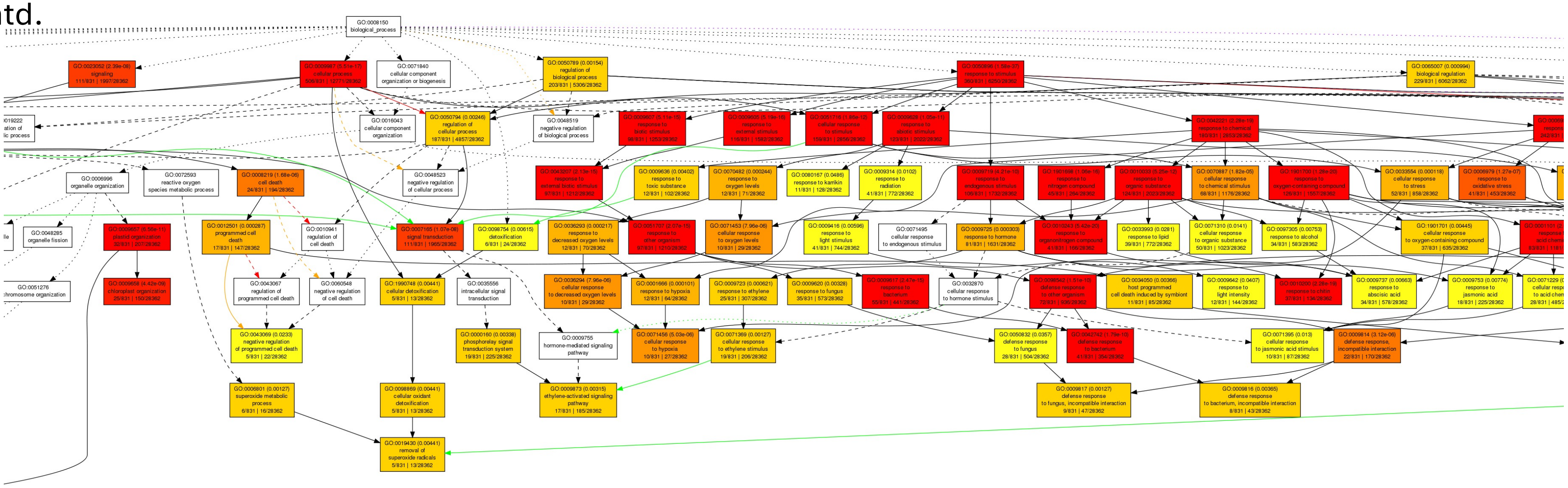

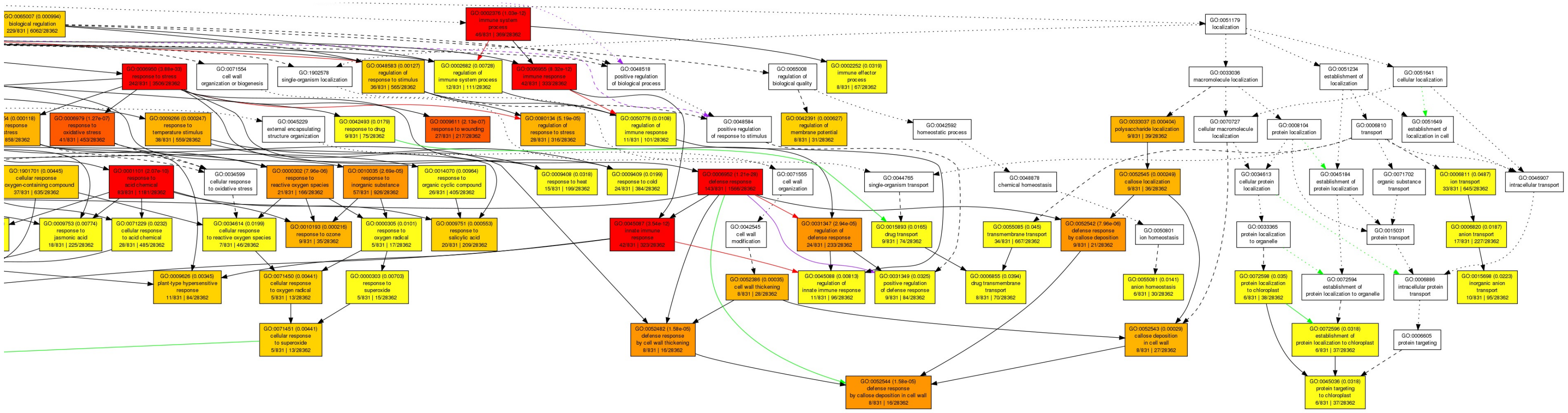

B

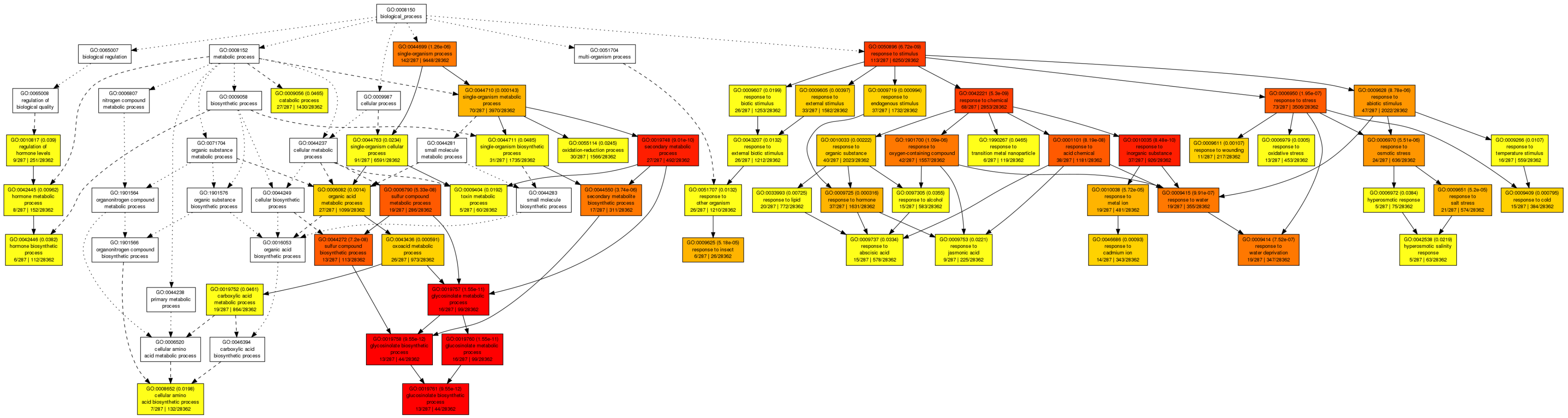
GO:0008150 biological_process
GO:0065007 biological regulation
GO:0008152 metabolic process
GO:0044699 (1.26e-06) single-organism process 142/287 | 9448/28362
GO:0051704 multi-organism process
GO:0050896 (6.72e-09) response to stimulus 113/287 | 6250/28362
GO:0065008 regulation of biological quality
GO:0006807 nitrogen compound metabolic process
GO:0009058 biosynthetic process
GO:0009056 (0.0465) catabolic process 27/287 | 1430/28362
GO:0009987 cellular process
GO:0044710 (0.000143) single-organism metabolic process 70/287 | 3970/28362
GO:0009607 (0.0199) response to biotic stimulus 26/287 | 1253/28362
GO:0009605 (0.00397) response to external stimulus 33/287 | 1582/28362
GO:0009719 (0.000994) response to endogenous stimulus 37/287 | 1732/28362
GO:0042221 (5.3e-09) response to chemical 68/287 | 2853/28362
GO:0006950 (1.95e-07) response to stress 73/287 | 3506/28362
GO:0009628 (8.78e-06) response to abiotic stimulus 47/287 | 2022/28362
GO:0010817 (0.039) regulation of hormone levels 9/287 | 251/28362
GO:0071704 organic substance metabolic process
GO:0044237 cellular metabolic process
GO:0044763 (0.0234) single-organism cellular process 91/287 | 6591/28362
GO:0044281 small molecule metabolic process
GO:0044711 (0.0485) single-organism biosynthetic process 31/287 | 1735/28362
GO:0055114 (0.0245) oxidation-reduction process 30/287 | 1566/28362
GO:0019748 (9.01e-10) secondary metabolic process 27/287 | 492/28362
GO:0043207 (0.0132) response to external biotic stimulus 26/287 | 1212/28362
GO:0010033 (0.00222) response to organic substance 40/287 | 2023/28362
GO:1901700 (1.09e-06) response to oxygen-containing compound 42/287 | 1557/28362
GO:1990267 (0.0465) response to transition metal nanoparticle 6/287 | 119/28362
GO:0001101 (8.19e-08) response to acid chemical 38/287 | 1181/28362
GO:0010035 (8.48e-10) response to inorganic substance 37/287 | 926/28362
GO:0009611 (0.00107) response to wounding 11/287 | 217/28362
GO:0006979 (0.0305) response to oxidative stress 13/287 | 453/28362
GO:0006970 (5.51e-06) response to osmotic stress 24/287 | 636/28362
GO:0009266 (0.0107) response to temperature stimulus 16/287 | 559/28362
GO:0042445 (0.00962) hormone metabolic process 8/287 | 152/28362
GO:1901564 organonitrogen compound metabolic process
GO:1901576 organic substance biosynthetic process
GO:0044249 cellular biosynthetic process
GO:0006082 (0.0014) organic acid metabolic process 27/287 | 1099/28362
GO:0006790 (5.33e-08) sulfur compound metabolic process 19/287 | 286/28362
GO:0009404 (0.0192) toxin metabolic process 5/287 | 60/28362
GO:0044283 small molecule biosynthetic process
GO:0044550 (3.74e-06) secondary metabolite biosynthetic process 17/287 | 311/28362
GO:0051707 (0.0132) response to other organism 26/287 | 1210/28362
GO:0033993 (0.00725) response to lipid 20/287 | 772/28362
GO:0009725 (0.000316) response to hormone 37/287 | 1631/28362
GO:0097305 (0.0355) response to alcohol 15/287 | 583/28362
GO:0010038 (5.72e-05) response to metal ion 19/287 | 481/28362
GO:0009415 (9.91e-07) response to water 19/287 | 355/28362
GO:0006972 (0.0384) hyperosmotic response 5/287 | 75/28362
GO:0009651 (5.2e-05) response to salt stress 21/287 | 574/28362
GO:0009409 (0.000795) response to cold 15/287 | 384/28362
GO:0042446 (0.0382) hormone biosynthetic process 6/287 | 112/28362
GO:1901566 organonitrogen compound biosynthetic process
GO:0016053 organic acid biosynthetic process
GO:0044272 (7.2e-08) sulfur compound biosynthetic process 13/287 | 113/28362
GO:0043436 (0.000591) oxoacid metabolic process 26/287 | 973/28362
GO:0009625 (5.18e-05) response to insect 6/287 | 26/28362
GO:0009737 (0.0334) response to abscisic acid 15/287 | 578/28362
GO:0009753 (0.0221) response to jasmonic acid 9/287 | 225/28362
GO:0046686 (0.00093) response to cadmium ion 14/287 | 343/28362
GO:0009414 (7.52e-07) response to water deprivation 19/287 | 347/28362
GO:0042538 (0.0219) hyperosmotic salinity response 5/287 | 63/28362
GO:0044238 primary metabolic process
GO:0019752 (0.0461) carboxylic acid metabolic process 19/287 | 864/28362
GO:0019757 (1.55e-11) glycosinolate metabolic process 16/287 | 99/28362
GO:0006520 cellular amino acid metabolic process
GO:0046394 carboxylic acid biosynthetic process
GO:0019758 (9.55e-12) glycosinolate biosynthetic process 13/287 | 44/28362
GO:0019760 (1.55e-11) glucosinolate metabolic process 16/287 | 99/28362
GO:0008652 (0.0198) cellular amino acid biosynthetic process 7/287 | 132/28362
GO:0019761 (9.55e-12) glucosinolate biosynthetic process 13/287 | 44/28362

C

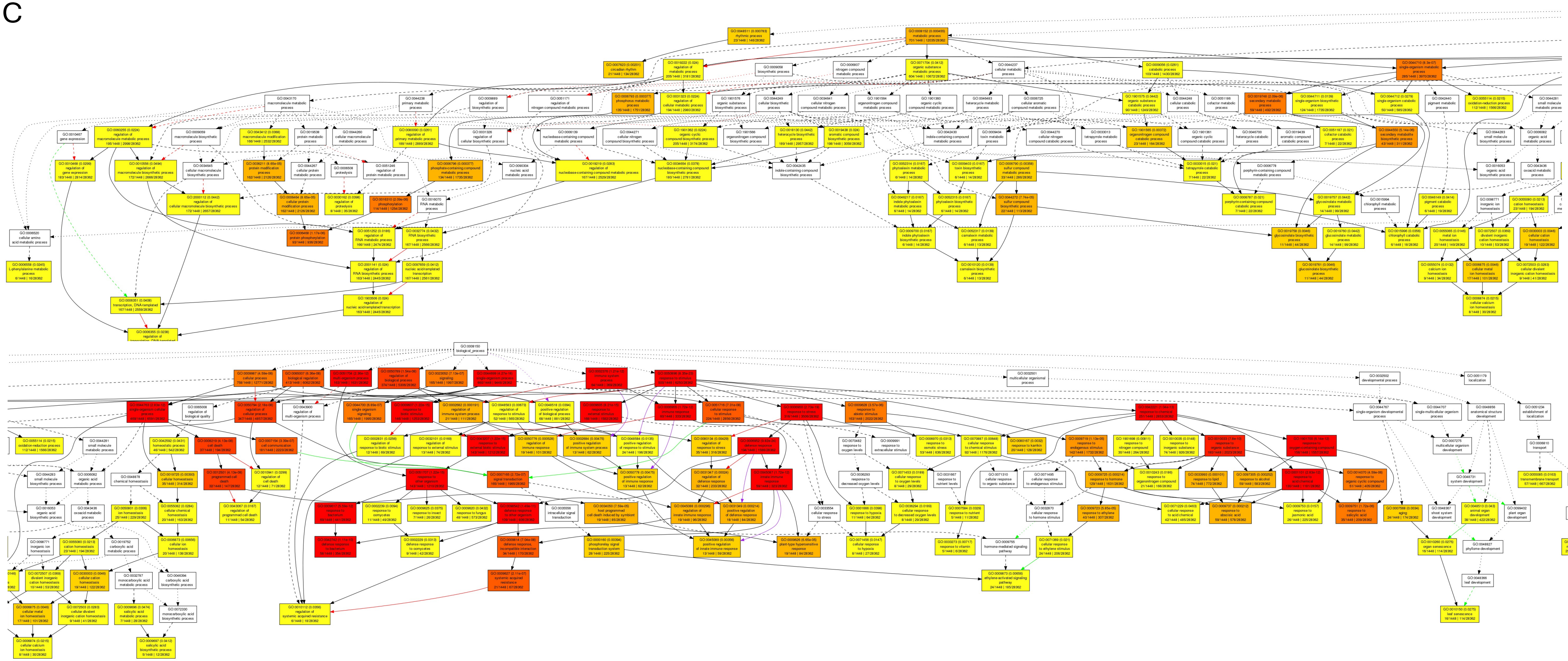

D

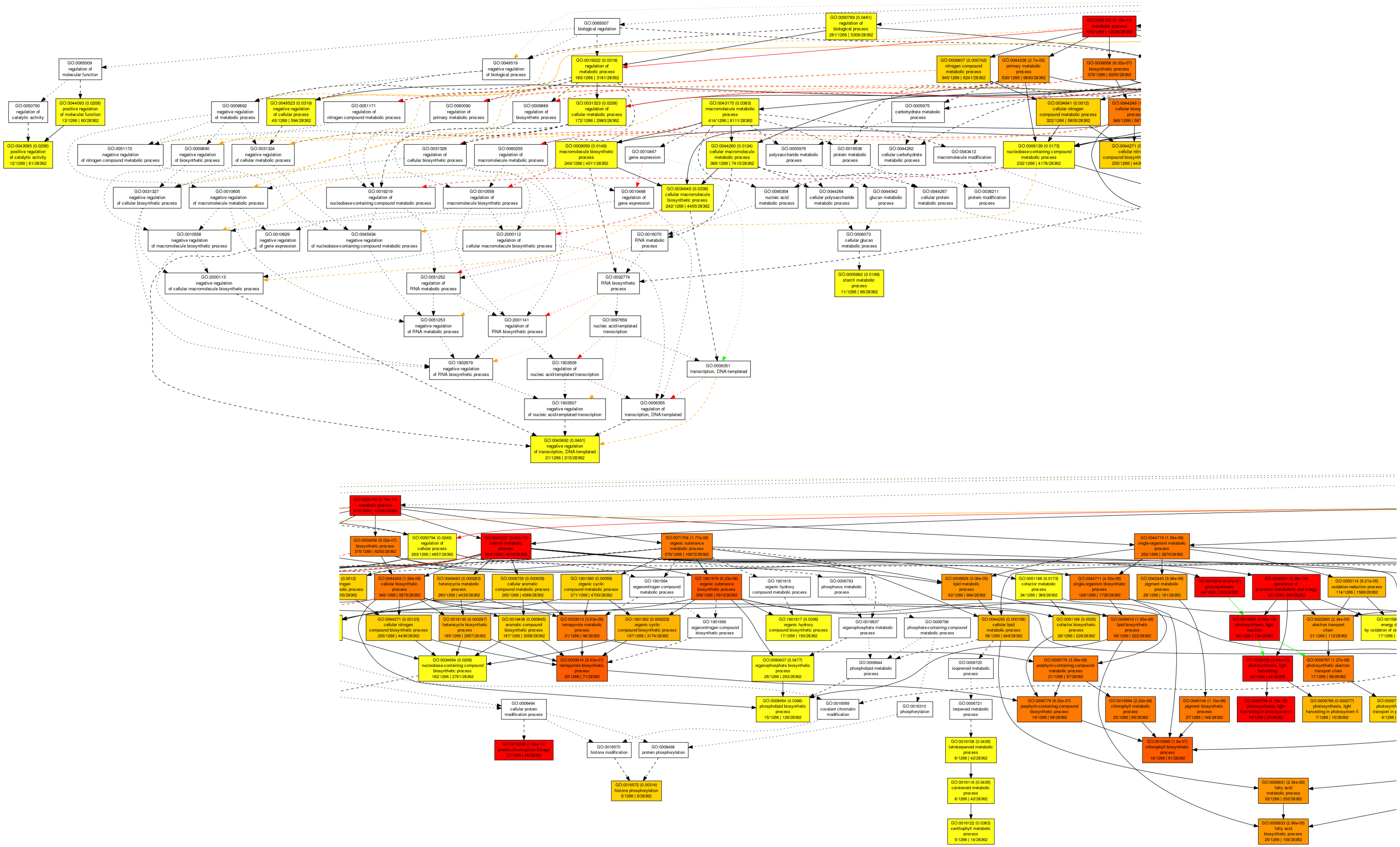

### D contd

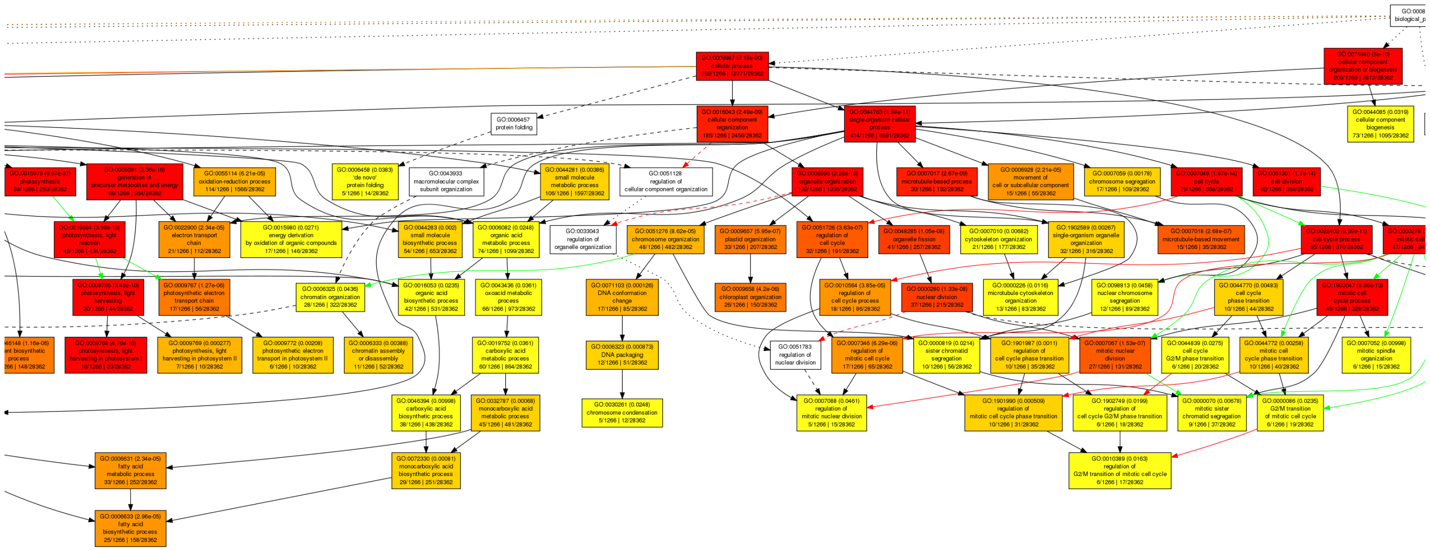

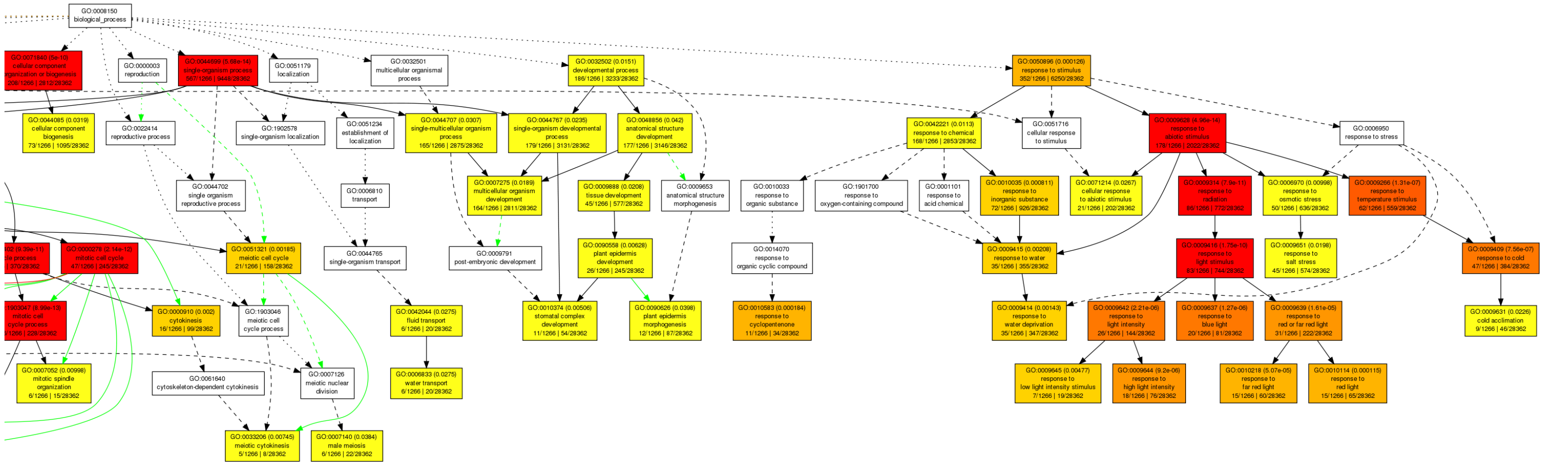

**Figure S2. RNA-seq and GO analysis of genes differentially in *pls* mutant and PLSOx seedlings at 7 d. post-germination compared to wildtype.**

(**A-D**) RNA-seq was carried out on RNA extracted from three biological replicate samples each of 7 day-old seedlings of wildtype, *pls* mutant and PLS over-expressing (PLSOx) transgenics. Bioinformatics and gene ontology (GO) analysis (AgriGO, www.bioinfo.cau.edu.cn/agriGO/) identified genes up-regulated (**A**) and down-regulated (**B**) in *pls* mutant seedlings compared to wildtype; and up-regulated (**C**) and down-regulated (**D**) in PLSOx seedlings compared to wild-type.

A

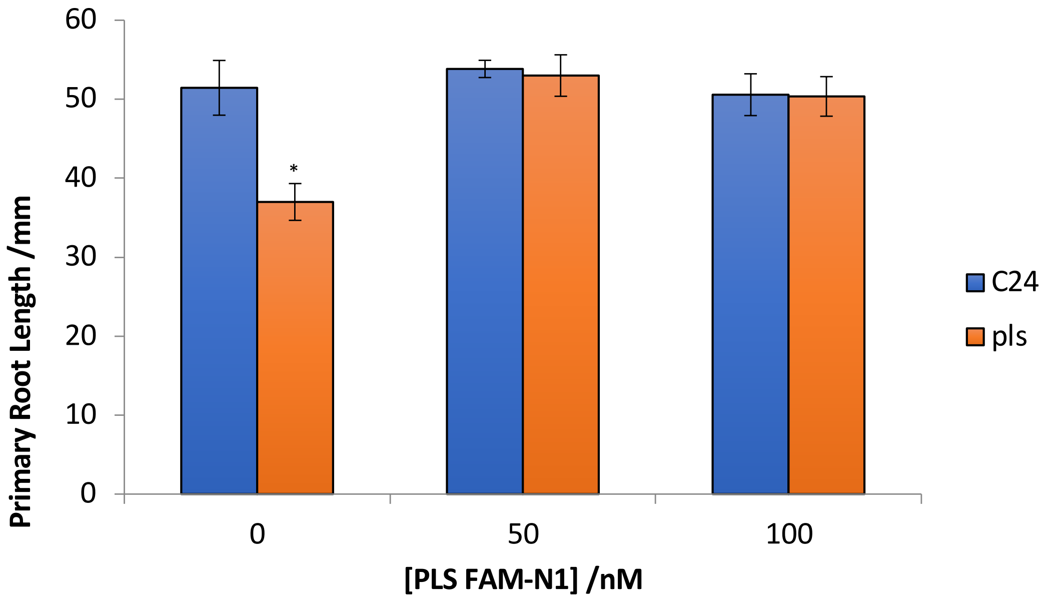


B

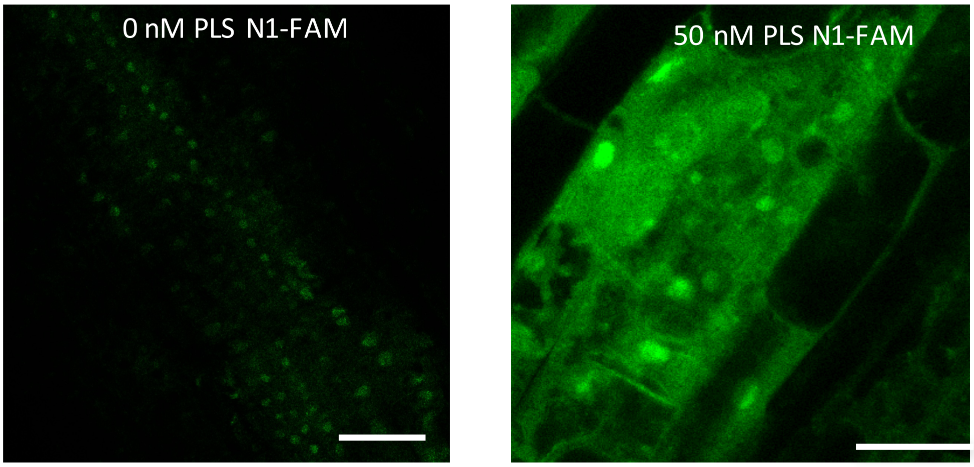


**Figure S3. Synthetic PLS peptide is taken up and rescues root development in *pls* seedlings.**

**(A) Effect of fluorescently tagged (5-FAM) PLS (N1) peptide on primary root length of *Arabidopsis*.**

Wildtype (C24, blue bars) and *pls* mutant (red bars) seedlings were grown hydroponically in the presence of 0, 50 or 100 nM peptide for 10 days. *: $P < 0.05$, $t$-test between C24 and *pls*, n = 15. Bars show ± standard errors of the mean, n = 15.

**(B) 5-FAM-PLS(N1) is taken up by *Arabidopsis* roots.** Confocal images of wildtype *Arabidopsis* roots (elongation zone) grown hydroponically for 10 days in the absence of peptide (left panel) and in the presence of 50 nM peptide (right panel). Scale bars = 10 μm.

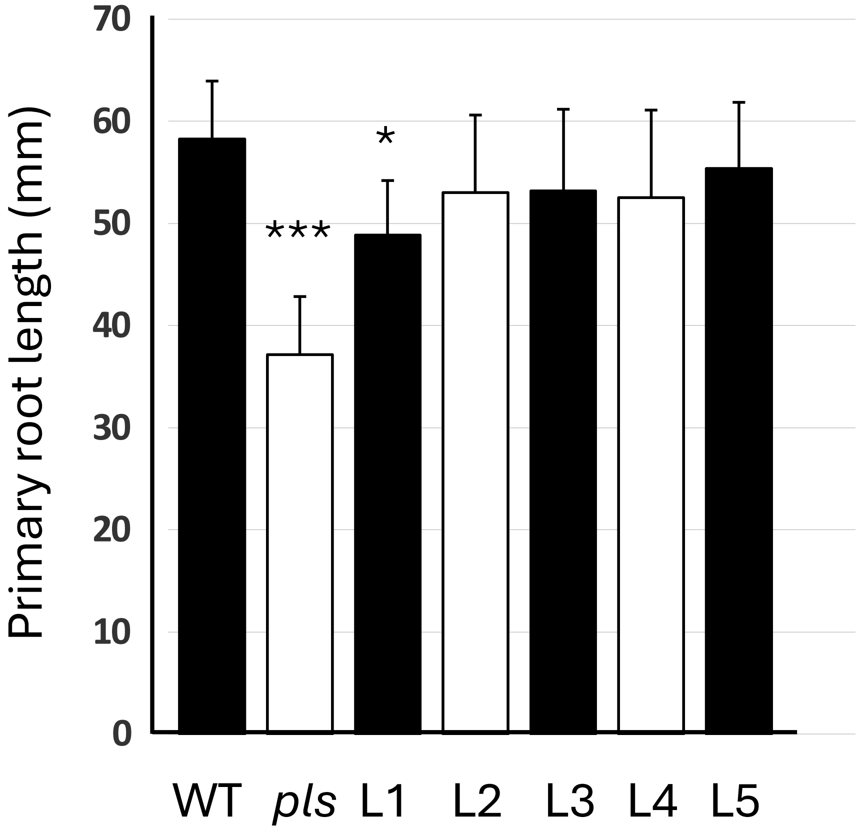


**Fig. S4. The *proPLS::PLS:GFP* gene fusion complements the *pls* mutant.** Primary root length of seedlings of wild type (WT), *pls* mutant (*pls*), and five independent transgenic lines (L1-L5) expressing*pro::PLS:PLS:GFP in the* pls mutant background. Seedlings were grown for 10 days on solid half MS10 agar medium. ***: *P* <0.0001, * : *P* <0.01 *t*-test between WT (C24) and *pls* or transgenic line, n = 13, bars represent mean values ± SD. No significant difference between WT and Lines 2-5.

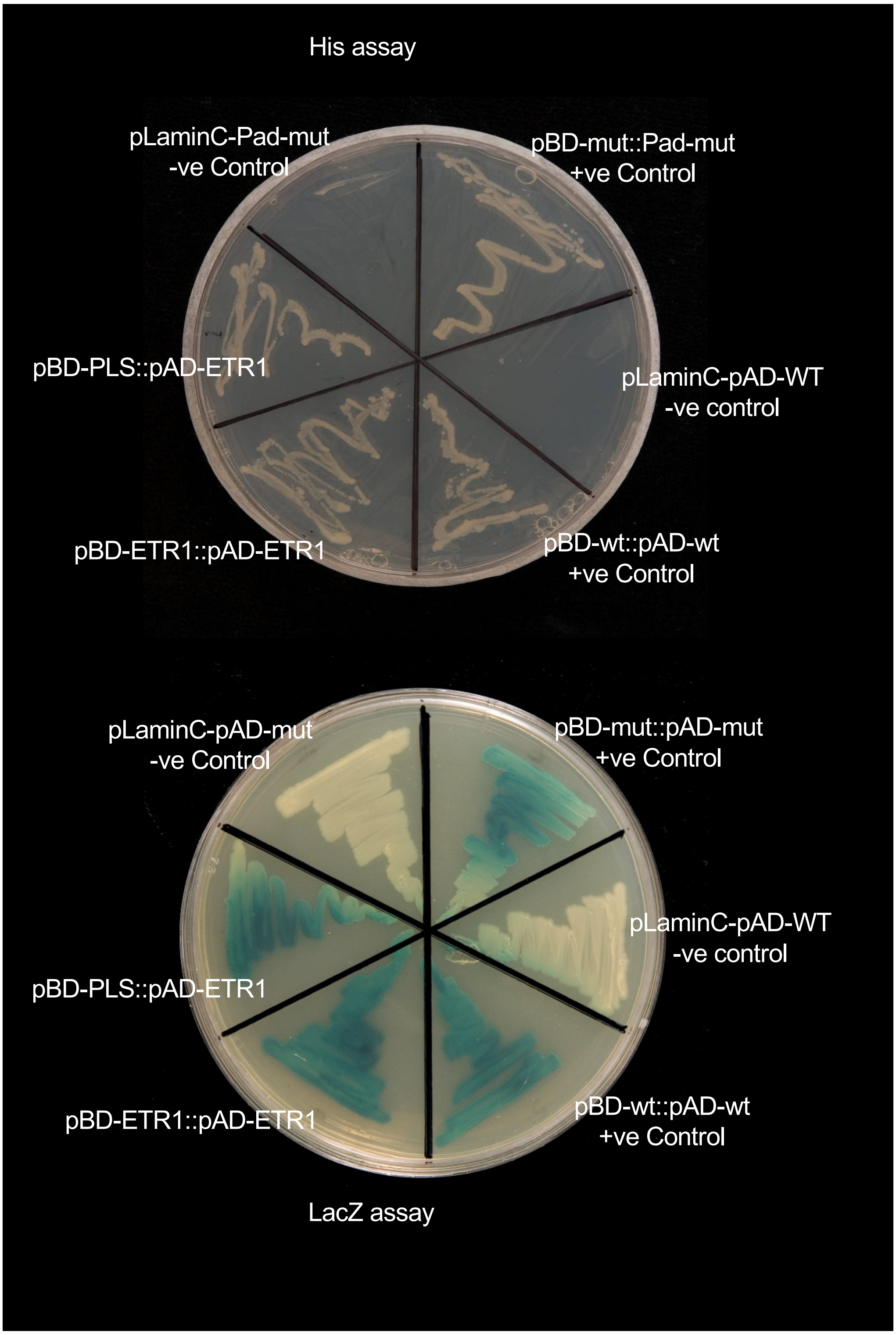


**Figure S5. PLS interacts with ETR1 in yeast.**

Yeast 2-hybrid assays showing and ETR1 interacts with PLS and with ETR1, in both His3 (upper plate) and LacZ (lower plate) reporter assays. Positive controls (pBD-wt::pAD-wt and pBD-mut::pAD-mut) and negative controls (pLaminC::pAD-mut and pLaminC::Pad-wt; pBD-PLS::pAD-ETR1 and pBD-ETR1::pAD-ETR1) are shown for each.

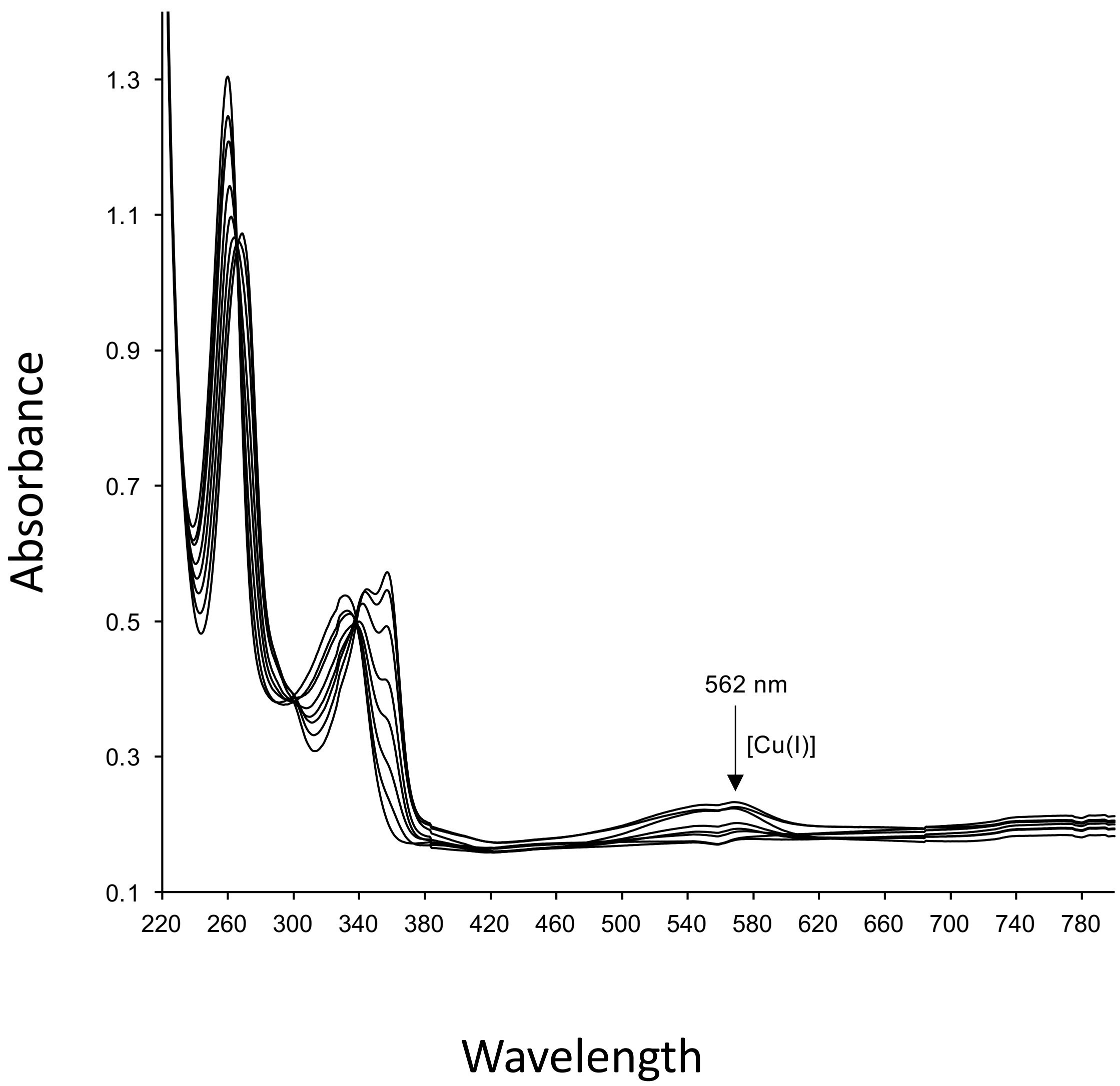


**Fig. S6. Full data representation from spectrum shown in Fig. 4D.** Titration of BCA with Cu(I) as in Figure 4D, showing additional copper-dependent features at 562 nm.

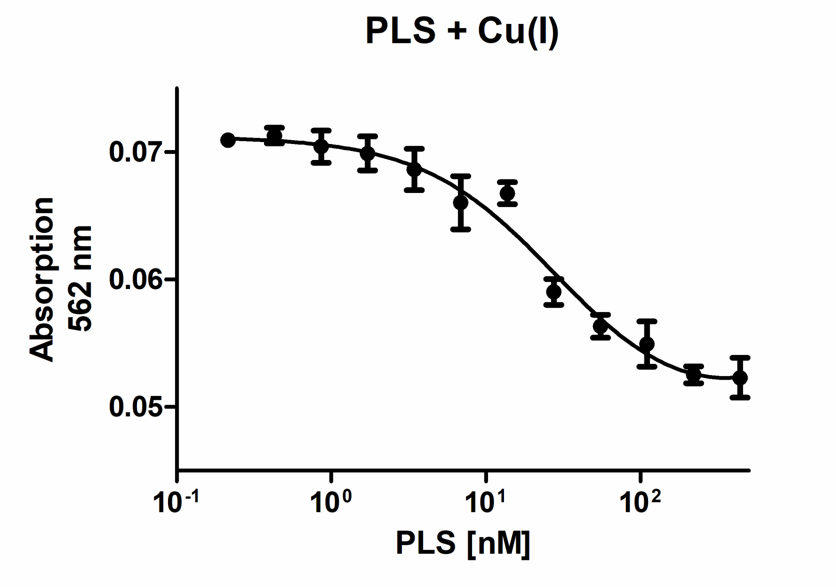


**Fig. S7. Titration of PLS in the presence of $BCA_2$-Cu(I)-Complex.**
A solution of PLS in 50 µl buffer (50 mM HEPES, 150 mM NaCl, pH 7.6) was serially diluted from 496 µM to 214 nM. After the addition of 50 µl $BCA_2$-Cu(I) buffer (50 mM HEPES, 150 mM NaCl, 20 mM ascorbate, 125 µM BCA, 50 µM CuCl, pH 7.6) the absorbance at 562 nm was measured. Discoloration of the pink $BCA_2$-Cu(I) complex visible in the decrease of absorption at higher peptide concentration indicates that PLS is able to effectively compete and remove copper from the high affinity copper chelator BCA.

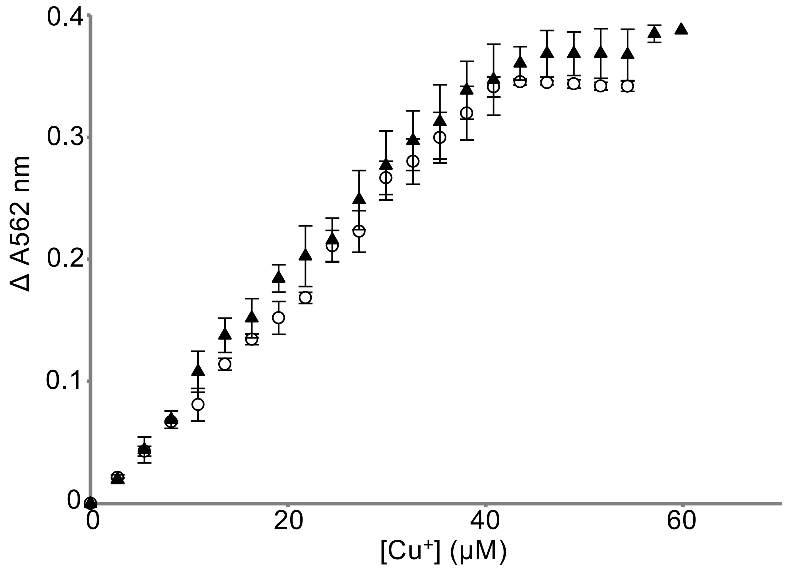


**Fig. S8. PLS requires C6S and C17S cysteine residues for copper binding.** BCA absorbance in the presence of PLS FL C6S, C17S ([BCA] = 93.3 μM, [PLS FL C6S, C17S] = 40 μM, closed triangles) or an equivalent volume (to PLS FL C6S, C17S) of DMSO ([BCA] = 87.4, open circles). Values represent means of three independent experiments with standard deviation.

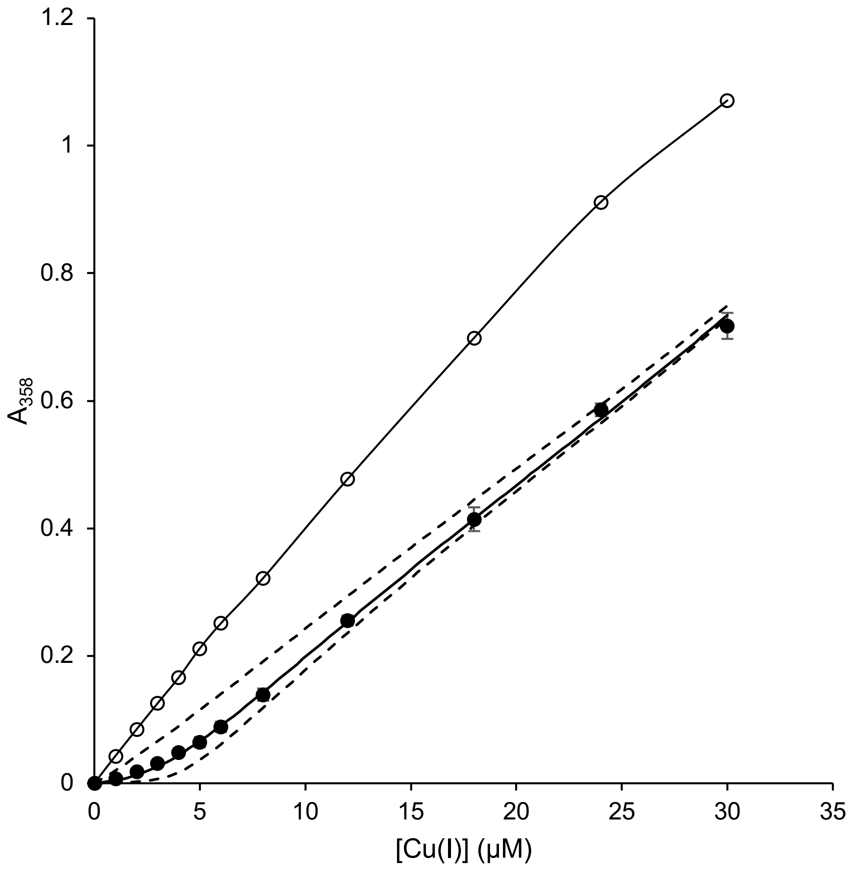


**Figure S9. Complete data set for Figure 4F.**

$A_{358\ nm}$ binding isotherm of BCA (50 µM) in the presence and absence of 10 µM MBP-PLS (filled and empty circles respectively) upon titration with $CuSO_4$ (recorded anaerobically at pH 7.0) in the presence of 1 mM $NH_2OH$. Dynafit model (solid line) describes Cu(I)-binding of MBP-PLS when modelled as forming a 2:1 complex with a $\beta_2$ value of 3.79 (±1.5) $x10^{19}$ $M^{-2}$ and one additional weak binding event. Dotted lines above and below represent the same model when the affinity is simulated 10x weaker or tighter, respectively (n=3, ±SD).

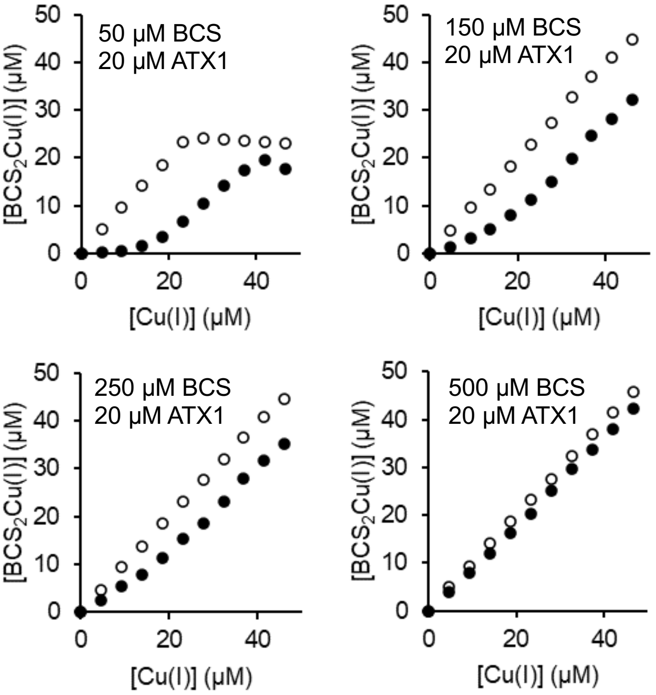


**Figure S10. Determination of the Cu(I) binding affinity of the *Arabidopsis* ATX1.**

Binding isotherms at $A_{483nm}$ when BCS (50, 150, 250 or 500 µM) in the presence and absence of 20 µM ATX1 (filled and empty circles, respectively), is titrated with Cu(I) under anaerobic conditions. Using the observed Cu(I):ATX1 stoichiometry of 1:1 (Fig. 4G, inset), affinities of 5.79 x$10^{-18}$, 4.83 x$10^{-18}$, 5.10 x$10^{-18}$, 6.14 x$10^{-18}$ respectively, with a mean of 5.47 (±0.6) x$10^{-18}$ M, were calculated.

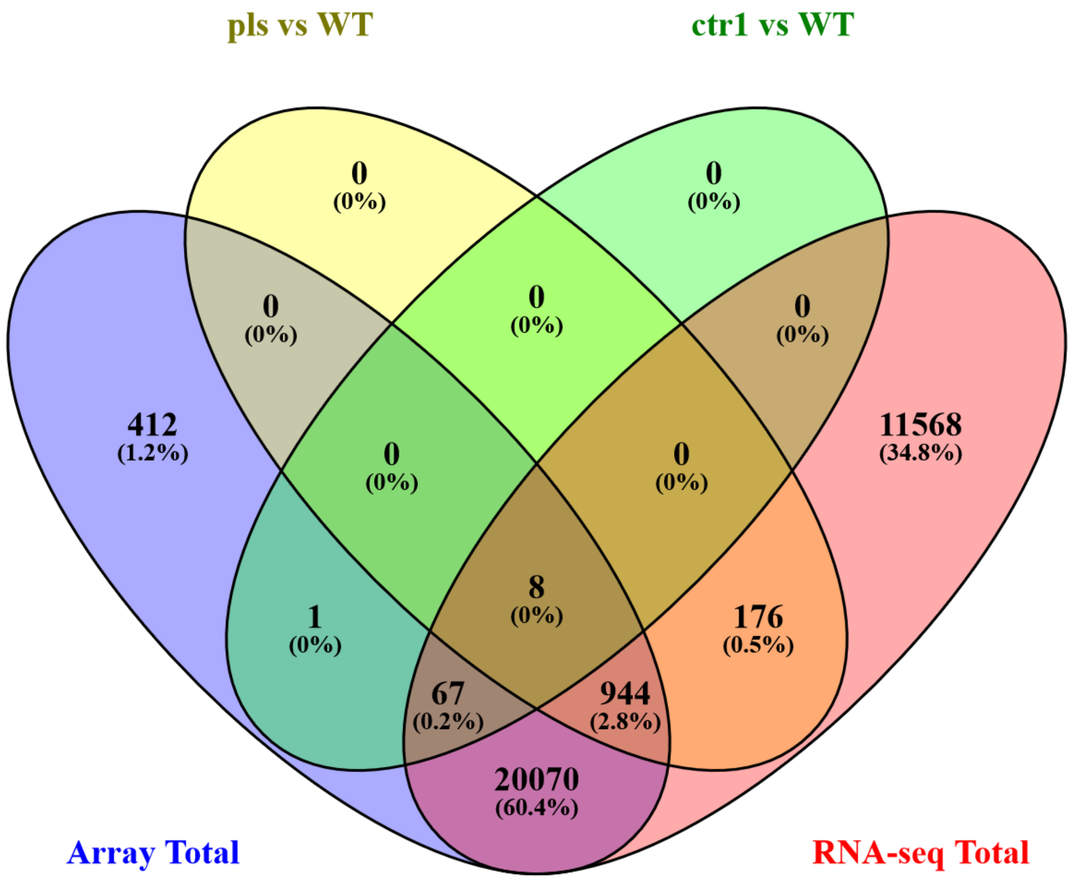


**Fig. S11 Overlap of DEGs (compared to respective wild types) between the *ctr1* and *pls* mutants.**

Venn diagram generated from DEG lists generated by Brodersen *et al*. (2006) *Plant J*. **47**, 532-546, Table S2 for *ctr1 vs.* wild type, and data from Table S1 (this paper) for *pls vs.* wild type. Overlap between the CTR1- and PLS-regulated genes is significant. P value of Fisher's exact test (alternative='greater') = 0.01959. 21089 genes are analysed in both experiments, of which the proportion of CTR1-regulated genes is 8+67/21089=0.36%. The proportion of CTR1- regulated genes in PLS-regulated genes is increased to 8/(8+944)=0.84%.

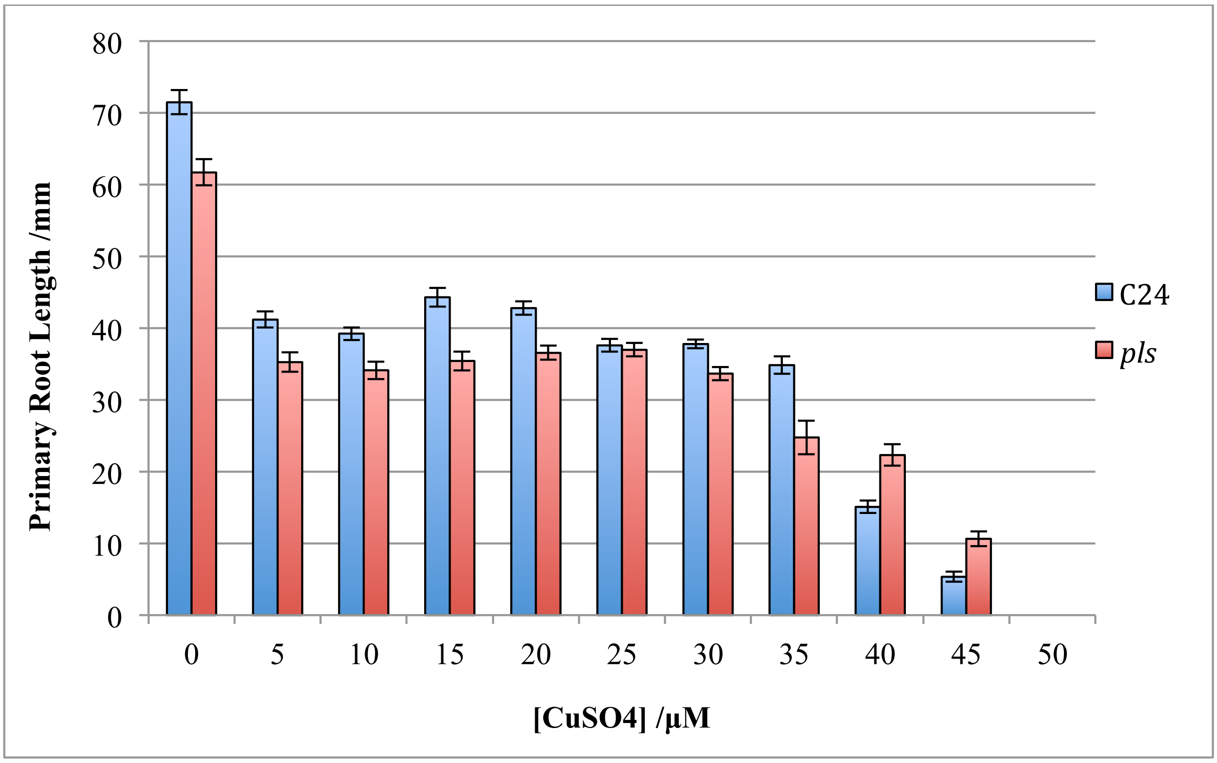


**Fig. S12. The *pls* mutant primary root is not hypersensitive to exogenous copper.**

Seedlings were grown for 12 days in liquid ½ MS10 media supplemented with $CuSO_4$. Root length was measured after 12 days using ImageJ. Error bars show ± 1 standard error. red bars represent *pls* mutant, blue bars represent wild type.

**Figure S13. The link between auxin and ethylene signalling effects on *PLS* transcription, and downstream effects on ethylene responses (a-d).**

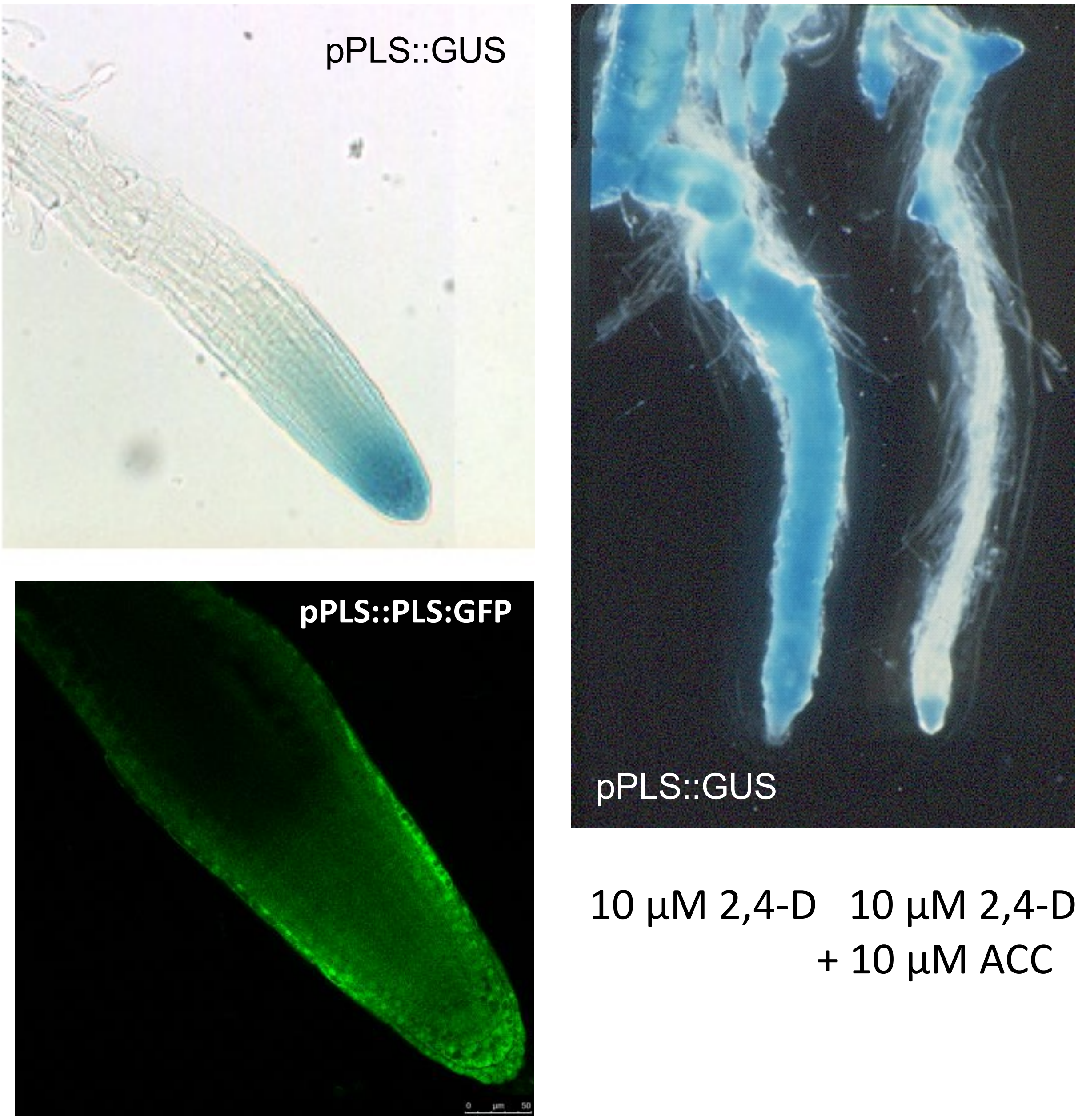


**(A) Antagonism between auxin and ethylene on PLS expression.**
Auxin positively regulates *PLS* gene transcription, while ACC (ethylene precursor) treatment antagonizes the inductive effects of auxin. Left upper panel: pPLS::GUS expression in the Arabidopsis root tip under standard growth conditions. Left lower panel: pPLS::PLS:GFP expression in the Arabidopsis root tip under standard growth conditions. Right panel: pPLS::GUS expression in root tips treated with 10 µM 2,4-D, showing ectopic expression throughout the root (left); and pPLS::GUS expression in root tips treated with 10 µM 2,4-D plus 10 µm ACC, showing the antagonism of the inductive effect of 2,4-D on PLS::GUS expression.

B

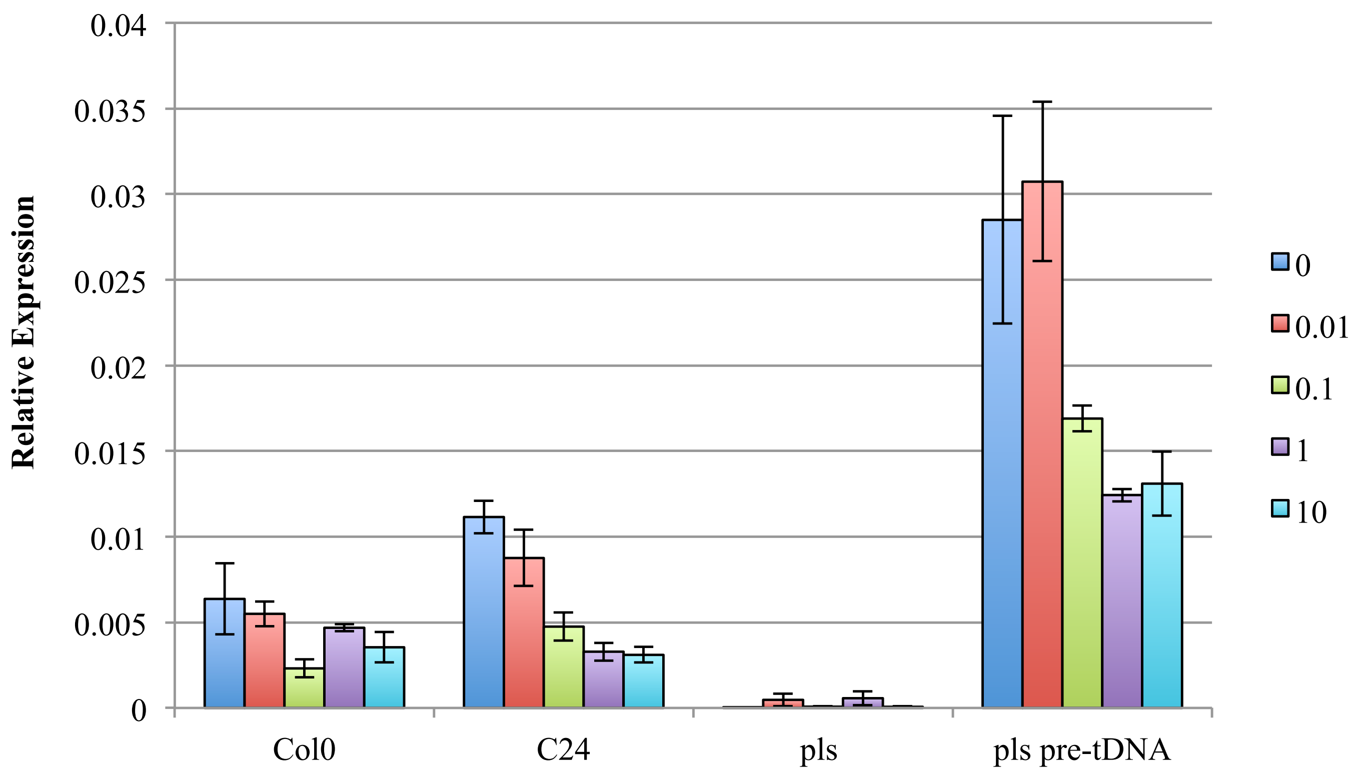


**(B) *PLS* gene expression under ACC treatment.**

Col-0, C24 and *pls* seedlings were treated with 0, 0.01, 0.1, 1 or 10 μM ACC for seven d.a.g and quantitative PCR was carried out on the resulting cDNA. *PLS* transcript levels were measured alongside a reference gene transcript *PP2C*. Comparison is made between the *PLS* expression in Col-0 and C24 wild types and the *pls* mutant, in which the reverse qPCR primer is located beyond the tDNA insertion in the *pls* mutant. In the fourth set of bars, a second set of *PLS* primers were used, located before the tDNA insertion, to detect a truncated and inactive *PLS* transcript. Error bars show ±1 standard error.

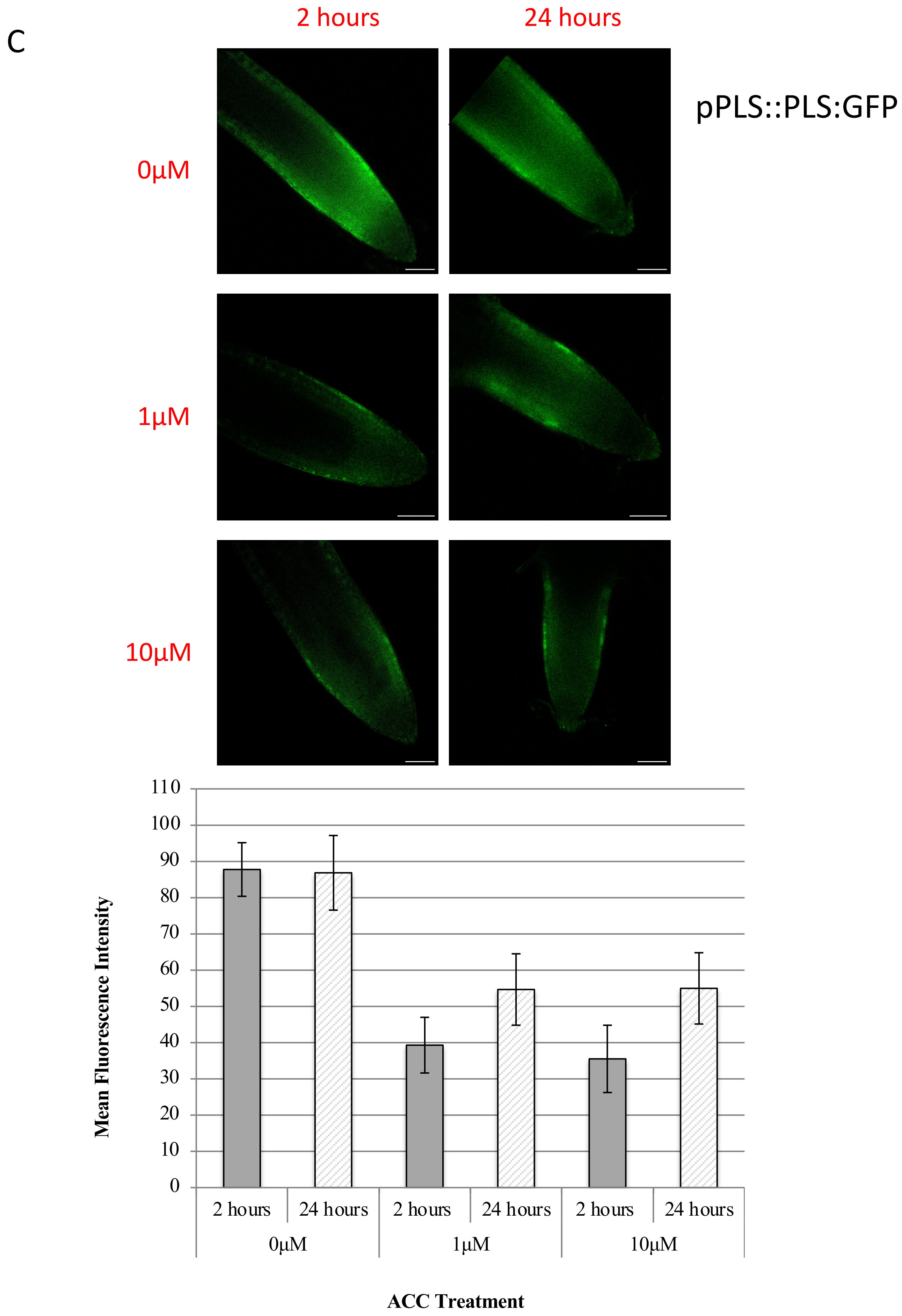
C
2 hours
24 hours
pPLS::PLS:GFP
0µM
1µM
10µM
Mean Fluorescence Intensity
0
10
20
30
40
50
60
70
80
90
100
110
2 hours
24 hours
2 hours
24 hours
2 hours
24 hours
0µM
1µM
10µM
ACC Treatment

**(C) Expression of the PLS-GFP fusion protein is downregulated in the root tip after ethylene treatment.**

Upper panel: Transgenic seedlings were treated with 0, 1 or 10 µM ACC for either 2 or 24 hours and imaged by CLSM at seven d.a.g. Laser settings were maintained at 21% 488 nm 20 mW, 970 V gain. Scale bars = 50 µm.

Lower panel: Quantification of PLS-GFP fluorescence after ACC treatment, detected in seedlings expressing the *pPLS::PLS:GFP* construct. Seedling root tips at seven d.a.g were imaged by CLSM after treatment with 0, 1 or 10 µM ACC for either 2 or 24 hours. Fluorescence intensity in each root was measured using ImageJ and the mean intensity for each treatment was calculated. Error bars show ± 1 standard error. N ≥ 17.

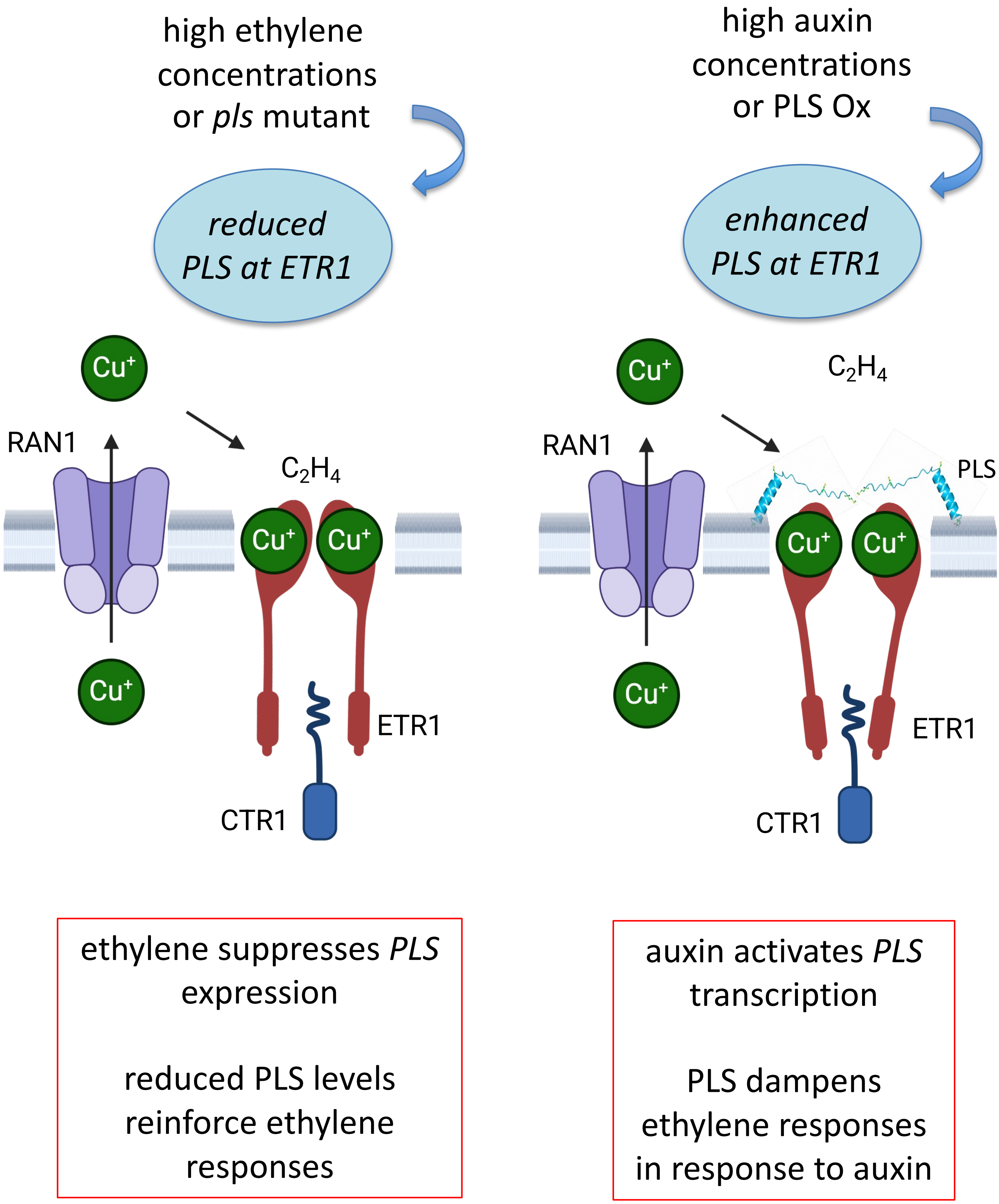

high ethylene concentrations or pls mutant
reduced PLS at ETR1
Cu+
RAN1
C2H4
Cu+
Cu+
Cu+
ETR1
CTR1
ethylene suppresses PLS expression
reduced PLS levels reinforce ethylene responses
high auxin concentrations or PLS Ox
enhanced PLS at ETR1
Cu+
C2H4
RAN1
PLS
Cu+
Cu+
Cu+
ETR1
CTR1
auxin activates PLS transcription
PLS dampens ethylene responses in response to auxin

**(D) Model for the regulation of ethylene responses by auxin and ethylene via effects on *PLS* expression.**

Left: Under high ethylene conditions or in the *pls* mutant, ETR1 and CTR1 are inactive, leading to enhanced ethylene responses. Right: Under high auxin conditions or in PLS-overexpressing plants, *PLS* transcription is relatively high, and PLS binds to the receptor to disrupt ethylene binding in an otherwise active receptor, so reducing ethylene responses. In for example a root tip with relatively high auxin concentrations, *PLS* expression would suppress ethylene responses to allow root growth; but under high ethylene conditions, root growth would be inhibited (as seen in the *pls* mutant).
