## Supplementary material for "POLARIS is a copper-binding peptide required for ethylene signalling control in *Arabidopsis*": Table S7 (from another version of this preprint: version 5, 2024-12-18)

**Table S7.**
**Primers for RT-qPCR**

| **Gene Forward/Reverse** | **Primer sequence 5'-3'** | **Tm (°C)** | **Product size (bp)** |
|---|---|---|---|
| *PLS* Forward | CAGAGAGAAAGAGAAGAGCACG | 58.5 | 130 |
| *PLS* Reverse | TAATTTCAGGCGAAGGTCCAT | 57.4 | |
| *PLS* pre-T-DNA Forward | GCAGTGTCTCACTGAAACATG | 57.5 | 132 |
| *PLS* pre-T-DNA Reverse | CAATGGATTTTAAAAAGTTTAAA CAATTTTGC | 58.3 | |
| *PP2C* Forward | AGCAGGGTGAGGATTTGGTG | 59.4 | 136 |
| *PP2C* Reverse | ATTCACCTGGCAAATCCGGT | 57.3 | |
| *ACTIN2* Forward | GGATCGGTGGTTCCATTCTTGC | 56 | 256 |
| *ACTIN2* Reverse | AGAGTTTGTCACACACAAGTGCA | 55 | |
